# Molecular signatures and cellular responses underlying hypoxic adaptation in sea turtles

**DOI:** 10.64898/2026.01.09.698755

**Authors:** B. Gabriela Arango, Giovanna Selleghin-Veiga, Diana Daniela Moreno-Santillán, David C. Ensminger, Juan Manuel Vázquez, Rebeca Tarvin, Mariana Nery, Céline A. Godard-Codding, José Pablo Vázquez-Medina

## Abstract

Hypoxia-inducible factors (HIFs) are transcriptional regulators that orchestrate the canonical response to low-oxygen tension in animal cells. Vertebrates possess three HIF-α isoforms, which arose from two gene duplication events of the ancestral HIF-1α gene. Here, we examined whether the HIF gene family (HIF-1α, HIF-2α, HIF-3α, and HIF-1αN inhibitor) shows evidence of positive selection in hypoxia-tolerant reptiles (Testudines), compared evolutionary patterns within the family, and assessed the transcriptional response to hypoxia in primary cells derived from a hypoxia-tolerant (*Caretta caretta*) and a non-tolerant (*Sceloporus occidentalis*) reptile. We found that HIF-1α, HIF-2α, and HIF-1αN are highly conserved across reptiles, whereas HIF-3α is under positive selection in Testudines. We also identified multiple novel regulatory motifs unique to Testudines. Transcriptional signatures of hypoxia exposure indicated stark differences between lizards and turtles. Whereas lizard cells exhibited a canonical response to hypoxia, characterized by enriched cell-survival pathways, sea turtle cells exhibit a robust, distinctive transcriptional response involving enriched pathways related to protein synthesis, quality maintenance, and mitochondrial integrity. Surprisingly, cis-regulatory element analysis did not show HIFs as key regulators of the transcriptional response in either species. Instead, TFDP1 in lizard cells and E2F1 in sea turtle cells emerged as potential key regulators. TFDP1 regulates the cell cycle, specifically DNA synthesis and cell cycle progression, while E2F regulates DNA-damage response, apoptosis, metabolism, and fatty acid biosynthesis. These results suggest that the reptilian response to hypoxia is shaped by transcriptional plasticity, while highlighting key regulatory mechanisms driving hypoxic adaptation in sea turtle cells. However, positive selection of HIF-3α and novel HIF motifs suggest a combined, but yet to be uncovered, contribution of regulatory and coding sequence evolutionary mechanisms shaping hypoxia tolerance in Testudines.

## INTRODUCTION

Hypoxia-inducible factors (HIFs) are key transcriptional regulators that orchestrate the canonical cellular response to low-oxygen availability (Majmundar *et al*., 2010). Under normoxia, HIF-α subunits are hydroxylated and targeted for degradation by Hypoxia Inducible Factor-inhibiting HIF (FIH, also known as HIF-1AN or HIF-1αN inhibitor) and prolyl hydroxylase domain (PHD)-containing proteins (Majmundar *et al*., 2010; Shi & Gilkes, 2025). A drop in oxygen tension prevents HIF degradation, allowing HIF-α subunits to bind p300/CBP coactivators and dimerize with HIF-1β or HIF-2β. These complexes accumulate rapidly, translocate to the nucleus, and bind hypoxia response elements (HREs)—specific DNA sequences that serve as binding sites for HIF proteins—inducing the expression of hundreds of genes involved in oxygen and metabolic homeostasis (Semenza, 2008; Shi & Gilkes, 2025). HIF-3α also functions as a transcription factor, acting both as a suppressor of HIF-1 and HIF-2 and as an activator of other target genes by binding to HREs (Yang *et al*., 2015). These central roles in oxygen sensing and response suggest that HIF genes may be subject to strong functional constraints or under positive selection.

Vertebrates’ three HIF-α isoforms arose from two gene duplication events of the ancestral HIF-1α gene. Phylogenetically, HIF-1α is more closely related to HIF-3α than to HIF-2α (Belato *et al*., 2024). Most evolutionary HIF studies have focused on fishes and mammals, including high-altitude-adapted humans. These studies show that HIF-1α is highly conserved across taxa, especially in species adapted to hypoxia, while HIF-2α shows substantial divergence (Rytkönen *et al*., 2011; Graham & Presnell, 2017; Zhu *et al*., 2018; Mandic *et al*., 2021). Reptiles have experienced wide fluctuations in atmospheric oxygen, especially during the Permian-Triassic period, when oxygen levels ranged from over 40% to below 15% (Mills *et al*., 2023). Testudines’ ancestors first appeared in the fossil record during this era and were likely subjected to strong selective pressures for hypoxia tolerance (Joyce, 2015), enabling modern testudines to occupy diverse ecological oxygen niches. Despite the remarkable ability of some reptiles, especially testudines, to withstand estivation, brumation, diving-induced hypoxia, and even anoxia, the evolutionary basis of reptile hypoxic tolerance remains poorly understood. Such extreme environmental constraints suggest that reptilian HIFs may have been subjected to selection in response to fluctuating oxygen availability.

Hypoxia-tolerant reptiles exhibit profound metabolic adaptations that enable survival under low oxygen availability, including post-translational protein modifications, differential gene expression, and constitutive activation of antioxidant systems that mitigate reoxygenation-induced oxidative stress (Storey, 1996; Bickler & Buck, 2007). Previous studies in cells derived from hypoxia-tolerant (sea turtles, which can remain submerged underwater for several hours) and non-tolerant reptiles (fence lizards) suggest functional differences in HIF-1α regulation. While both species stabilize HIF-1α under hypoxia, sea turtle cells show rapid nuclear accumulation, suggesting faster stabilization in the hypoxia-tolerant species. Intriguingly, sea turtle cells do not exhibit a canonical metabolic response to hypoxia. In contrast, lizard cells activate glycolytic metabolism upon hypoxic onset but fail to restart respiration upon reoxygenation (Arango *et al*., 2025). Despite this metabolic divergence, the faster nuclear accumulation of HIF-1α in sea turtle cells suggests that stabilization occurs rapidly. This raises the possibility that, although the molecular mechanism of HIF stabilization is conserved, the downstream cellular response has diverged. These findings suggest that while sea turtle HIFs may have evolved to respond rapidly to hypoxia, the cellular response to low oxygen tension might involve non-canonical regulators, as observed in other hypoxia-tolerant vertebrates, including deep-diving seals, in which HIF-1 activation is decoupled from canonical signaling pathways in vascular cells (Allen *et al*., 2024).

Here, we tested whether HIFs genes (HIF-1α, HIF-2α, HIF-3α, and HIF-1AN) show evidence of positive selection in reptiles, compared evolutionary patterns within the HIF gene family, and measured hypoxia-induced transcriptional shifts in cultured cells derived from a hypoxia-tolerant (loggerhead sea turtle, *Caretta caretta*) and non-tolerant reptile (western fence lizard, *Sceloporus occidentalis*). Given the conserved nature of the HIF gene family, we aimed to determine whether differences in hypoxia tolerance arise primarily from changes in regulatory elements rather than in sequence variation, and that HIFs function as the primary regulators of the hypoxic response in these species. We hypothesized that HIF genes in sea turtle cells have undergone positive selection and that their transcriptional response to hypoxia is more robust (i.e. greater number of DE genes) than that of lizard cells. Our results show species-specific transcriptional responses to hypoxia. Lizard cells exhibit a canonical response with enriched pathways for cell-survival. In contrast, sea turtle cells activate a broader transcriptional response with enriched pathways involving protein synthesis, quality control, and mitochondrial maintenance. Strikingly, these differences appear to be driven by regulatory plasticity with limited HIF involvement. Moreover, our analysis suggests that TFDP1/E2F1 are potential key regulators of the hypoxic response in cultured reptile cells.

## METHODS

### Motif characterization and phylogenetic inference of reptilian HIF genes

Orthologous genes were identified using OrthoFinder v3.0 (Emms & Kelly, 2019) for each of our 48 annotated reptilian genomes retrieved from NCBI database (**Table 1**). In addition, *Homo sapiens* and *Gallus gallus* were used as outgroups. Using *Homo sapiens* gene IDs for each of the know HIF genes (HIF-1α, HIF-2α, HIF-3α, and HIF-1αN), we identified Phylogenetic Hierachical Orthogroups (HOGs) and used those to obtain the coding sequences (CDS). After a first trimming, the number of sequences varied by gene: HIF-1α = 49 sequences, HIF-2α = 50, HIF-3α = 36, and HIF-1αN = 44 sequences (**Table 1**). We also recovered sequences from four unannotated species of sea turtles (*Eretmochelys imbricata, Lepidochelys kempii, Lepidochelys olivacea, Natator depressus)* using BLAST (Altschul *et al*., 1990) with *Caretta caretta* sequences as queries.

**Table 1.**
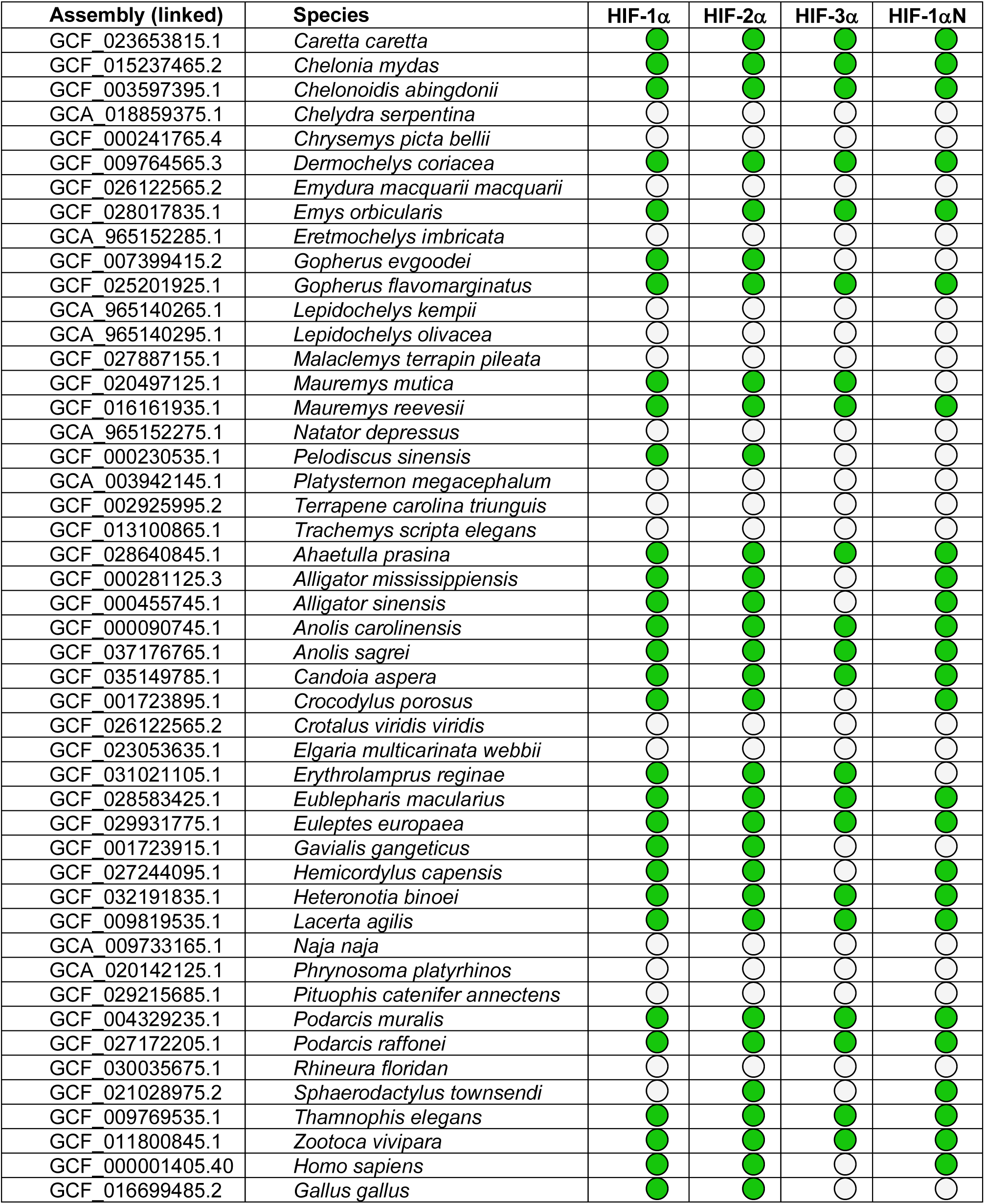
Species used in this study with genome assembly accession numbers and the presence of HIF genes. . The listed species correspond to those with publicly available full genome assemblies, retrieved from the NCBI database. Orthologous HIF genes (HIF-1α, HIF-2α, HIF-3α, and HIF-1αN) were identified using OrthoFinder across 48 annotated reptilian genomes. Gene presence was determined based on coding sequences (CDS) extracted from these assemblies. Green circles indicate CDS presence in the final alignment; white circles indicate absence. Additional species not included in this table lacked complete genome assemblies but were still represented in the HIF alignments based on publicly available CDS data.

To ensure intron-free alignments, CDS were manually curated against annotated reference HIF genes. For downstream analyses, CDS were initially aligned with MUSCLE v5.1 (Edgar, 2022) to construct phylogenetic trees. Alignments were then re-aligned using MAFFT v7.490 (Auto algorithm, BLOSUM62 matrix, gap open penalty of 1.53, and offset value of 0.123) (Katoh & Standley, 2013) to preserve codon triplets, and manually corrected to resolve misalignments. The final alignments were translated into proteins and used for selection analyses in *codeml*. CDS alignments were conducted in Geneious Prime v2025.1.2.

The regulatory regions—promoters were extracted from 1500 bases upstream of each gene transcript from all annotated reptile genomes. Promoter sequences were used to find five motifs per HIF gene using the MEME suite (Bailey *et al*., 2015). Motifs were compared against the JASPAR vertebrate and UniPROBE mouse databases using TomTom (Gupta *et al*., 2007), with a Pearson correlation coefficient and a threshold e-value < 0.05.

To validate alignment accuracy, we performed nucleotide or protein BLAST searches on 2-3 representative sequences from each alignment. Phylogenetic trees were reconstructed from promoters and CDS alignments using Maximum Likelihood (ML) with ModelFinder (Kalyaanamoorthy *et al*., 2017) and ultra-fast bootstrapping (1000 iterations) to assess nodal support (Hoang *et al*., 2018), implemented in IQ-TREE v1.6.2 (Nguyen *et al*., 2015). Final tree topologies were visualized using iTol v7.1 (Letunic & Bork, 2024).

### Analysis of positive selection

We assessed positive selection using codon-based branch-site models implemented in *codeml* from PAML v.3 (Yang, 2007). Coding sequence alignments and the corresponding species phylogeny were used to estimate the ratio of nonsynonymous to synonymous substitutions (ω = *d*N/*d*S), measuring the strength and mode of natural selection acting on the protein genes. A value of ω > 1 indicates positive selection (adaptive), ω = 1 neutral evolution, and ω < 1 negative selection (purifying).

We applied the branch-site model (model = 2), which allows ω to vary among codon sites and across phylogenetic branches. This model enables the detection of site-specific positive selection affecting a subset of codons along specified foreground lineages. In our analysis, we designated a subset of reptilian lineages as foreground branches in two tests: 1) Testudines vs. other reptiles, and 2) sea turtles vs. other reptiles. For each test, we used likelihood ratio tests (LRTs) to compare the alternative model (allowing ω > 1 on the foreground branches) against a null model in which ω was fixed at 1. Significance was calculated under a chi-square distribution with one degree of freedom.

### Cellular experiments

Primary dermal fibroblasts from sea turtles (*C. caretta*) and lizards (*S. occidentalis*) were derived, maintained, and cultured as described in our previous work (Webb *et al*., 2014; Arango *et al*., 2025). Five conditions were used for hypoxia and hypoxia/reoxygenation experiments (Arango *et al*., 2025). Briefly, for hypoxic exposure, cells were incubated at 0.1% O_2_, 5% CO_2_ for 1 or 24 hours in an InVivO_2_ physiological workstation (Baker Ruskinn). Control conditions (normoxia) were 21% O_2_ and 5% CO_2_. Experimental treatments induce robust HIF-1αaccumulation without causing cell death in both lizard and sea turtle cells (Arango *et al*., 2025). For hypoxia/reoxygenation experiments, cells were incubated for 1 hour or 24 hours at 0.1% O_2_ and 5% CO_2_, followed by 1-hour reoxygenation (room air, 21% O_2_). Each sample of 1 million cells per treatment (cell pools from five individuals per species) was run in triplicate.

### RNA extraction and RNA-seq

Control, hypoxic, and reoxygenated cells were rinsed in PBS and lysed in RLT buffer without beta-mercaptoethanol (BME). RNA was extracted using a RNeasy Plus Mini Kit (Qiagen #74136), following the manufacturer’s instructions. An on-column DNase digestion step was included to remove genomic DNA. RNA was quantified using a Nanodrop spectrophotometer (ThermoScientific). Library preparation and sequencing were conducted by Novogene, with intermediate quality control checks at each step to ensure sample integrity and optimal sequencing. RNA integrity was assessed using a Bioanalyzer, with all samples achieving an RIN > 9. mRNA was enriched using poly-T oligo-attached magnetic beads, fragmented, and reverse-transcribed to synthesize first- and second-strand cDNA. The library was ready after end repair, A-tailing, adapter ligation, size selection, amplification, and purification. Paired-end sequencing was conducted on an Illumina PE150 platform with a sequencing depth of ∼50M reads per sample, yielding 3.6 GB of coverage for the lizard cells and 2.7 GB for the sea turtle cells.

### RNA-seq data processing and transcriptome analysis

Raw reads were filtered to remove adapter sequences and reads containing more than 10% ambiguous bases or reads with over 50% of bases with a quality score ≤ 5. Reads were processed and cleaned using TrimGalore (Bolger *et al*., 2014). Quality was assessed using FastQC (v. 0.12.1). Reads were mapped to the loggerhead sea turtle chromosome reference genome (GCF_965140235.1; Yen et al., 2024) using STAR (Dobin *et al*., 2013). BRAKER (v. 3.0.8) was run using RNA-Seq and protein data with GeneMark-ETP. We use the ensembl metazoa protein database and RNA-seq raw reads from *S. occidentalis* (reference genome GCA_023333645.1) to train BRAKER (Accession Numbers: SRR29377057, SRR29377058, SRR29377059, SRR29377060, SRR29377061, SRR29377062, SRR29377063) (Stanke *et al*., 2006, 2008; Quinlan, 2014; Kovaka *et al*., 2019; Pertea & Pertea, 2020; Gabriel *et al*., 2021, 2024; Brůna *et al*., 2024). Unique reads with mapping quality > 90% were retained. Transcript abundance was quantified using RSEM (Li & Dewey, 2011).

Gene expression data across different treatment conditions were normalized using transcripts per million (TPM) and grouped into two clusters: (a: short-term hypoxia) control—as a reference, 1-hour hypoxia, and 1-hour reoxygenation, and (b: long-term hypoxia) control, 1-hour hypoxia, 24-hour hypoxia, and reoxygenation after 24 hours hypoxia to assess gene expression trajectories across hypoxia exposure time. Median z-scores were calculated and used to construct a hierarchical clustering matrix. Pairwise distances were computed using the Euclidean Distance, and clustering was performed using the complete linkage method, which measures the distance between clusters as the maximum pairwise distance between points in two clusters.

K-means clustering, an unsupervised learning method used to group similar expression patterns, was performed across k-values and evaluated using silhouette scores (1 to 50), to assess cluster cohesion. The optimal number of clusters (k=6) was determined based on an Elbow plot (**Supplementary Figure 1**). Clusters were visualized based on gene expression trajectories, with each line representing a gene’s median Z-score across treatments.

Differential expression analysis was conducted using EBSeq on gene-level normalized counts from the GeneMat output. Differentially expressed (DE) genes between control and treatment groups were identified using a 5% false discovery rate (FDR), based on the posterior probability of differential expression. Pairwise DE comparisons were used for subsequent enrichment analysis. Gene Set Enrichment Analysis was used to identify significantly enriched pathways within clusters based on Gene Ontology (GO) Biological Processes (gene and protein functions), Mouse Genome Informatics (MGI) Mammalian Phenotype (gene-phenotype associations in mouse), and Kyoto Encyclopedia of Genes and Genomes (KEGG) pathways (biological pathways and metabolic reactions) using the Reactome database v93. Pathway enrichment was conducted using the Enrichr-KG platform (Evangelista *et al*., 2023). *Cis*-regulatory element analysis and Functional Interaction Networks (FIN) were constructed in Cytoscape v3.10.3 (Janky *et al*., 2014) using genes DE in each condition.

## RESULTS

### Phylogenetic relationships, positive selection, and promoter divergence in reptilian HIF genes

We compared evolutionary patterns within the reptilian HIF gene family (HIF-1α, HIF-2α, HIF-3α and HIF-1α inhibitor), explored whether Testudine HIFs are under selection, and analyzed gene expression profiles in hypoxic cells derived from a hypoxia-tolerant (sea turtle) and a non-tolerant (lizard) reptile to test whether hypoxia tolerance arises from genetic sequence variation or regulatory expression patterns. Phylogenetic Hierarchical Orthogroups (HOGs) corresponding to each HIF gene revealed that most species possess a single gene copy. Gene duplications were rare, and no consistent patterns of evolutionary divergence were observed across species (**Figure 1**). HIF-3α was the only gene that showed evidence of gene loss in some crocodilian species and in *Gallus gallus*.

**Figure 1.**
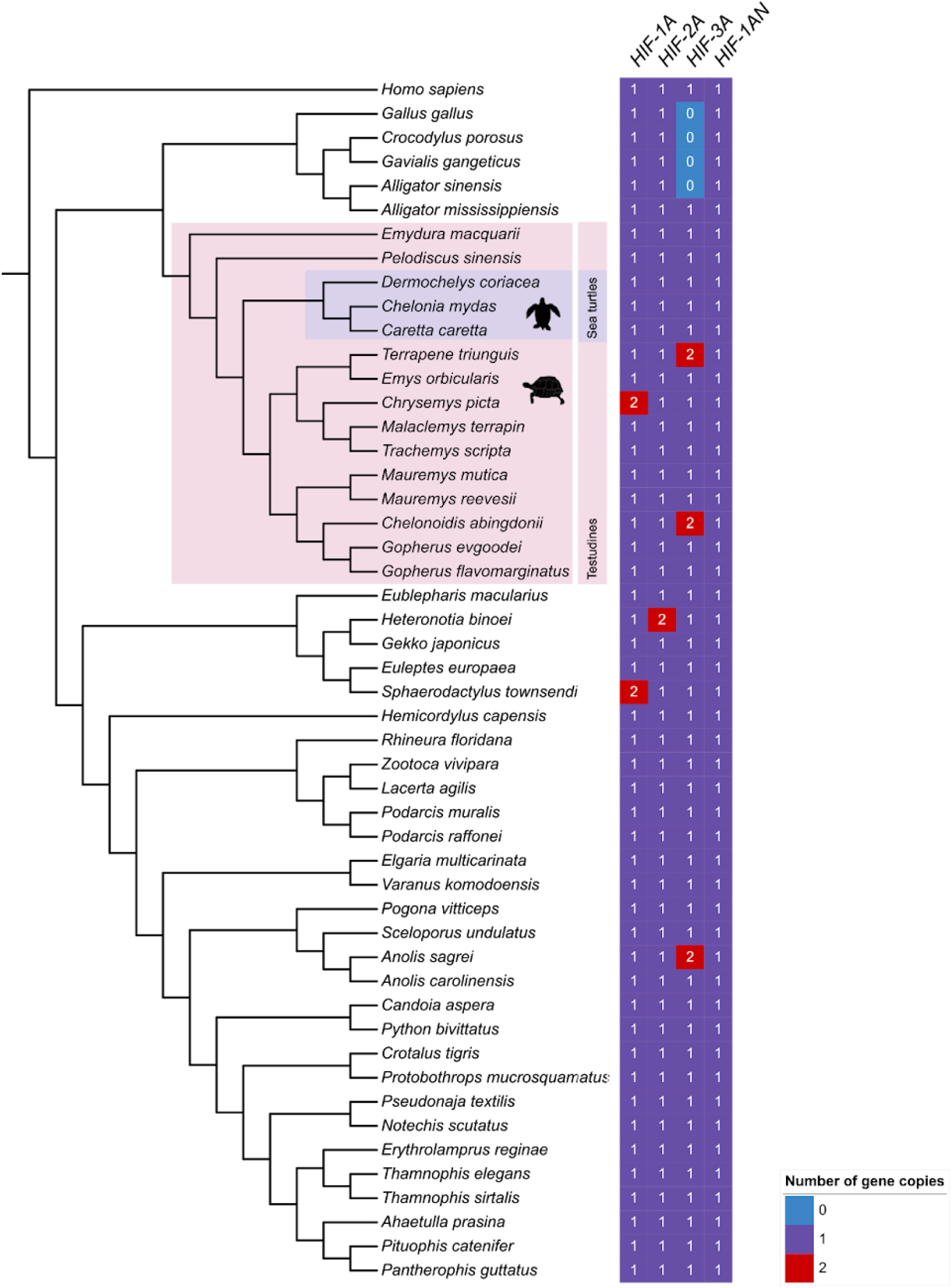
Reptile phylogeny and HIF gene copy number from OrthoFinder. Maximum likelihood tree based on a multiple sequence alignment of 2,239 single-copy genes, estimated using OrthoFinder. Heatmap shows gene copy number for each HIF gene (*HIF-1α, HIF-2α, HIF-3α*, and *HIF1AN*). Pink: Testudines (turtles, tortoises, and terrapins); Purple: Sea turtles.

Species tree reconstruction in OrthoFinder was based on a multiple sequence alignment of 2,239 single-copy genes and was rooted using *Homo sapiens* (**Figure 1**). The resulting topology closely matched the reptilian species tree in TimeTree (https://timetree.org/; **Supplementary Figure 2**), which also shows that Testudine ancestors emerged during the late Permian, a period of declining atmospheric oxygen that may have shaped the evolution of hypoxia tolerance in extant turtles (Mills *et al*., 2023). Based on this reconstructed phylogeny, we compared each gene tree generated by our HIF alignments. Similarly, HIF gene trees showed strong node support and there were no major differences between gene tree and species tree topologies, suggesting that HIF phylogeny is highly conserved among reptiles, including Testudines (**Figure 2a-d**). When analyzing all HIF genes, HIF-1αN is the sister lineage to a clade containing HIF-3α, HIF-1α, and HIF-2α (**Figure 3**). HIF-1α and HIF-2α share a more recent common ancestor with each other than to HIF-3α.

**Figure 2.**
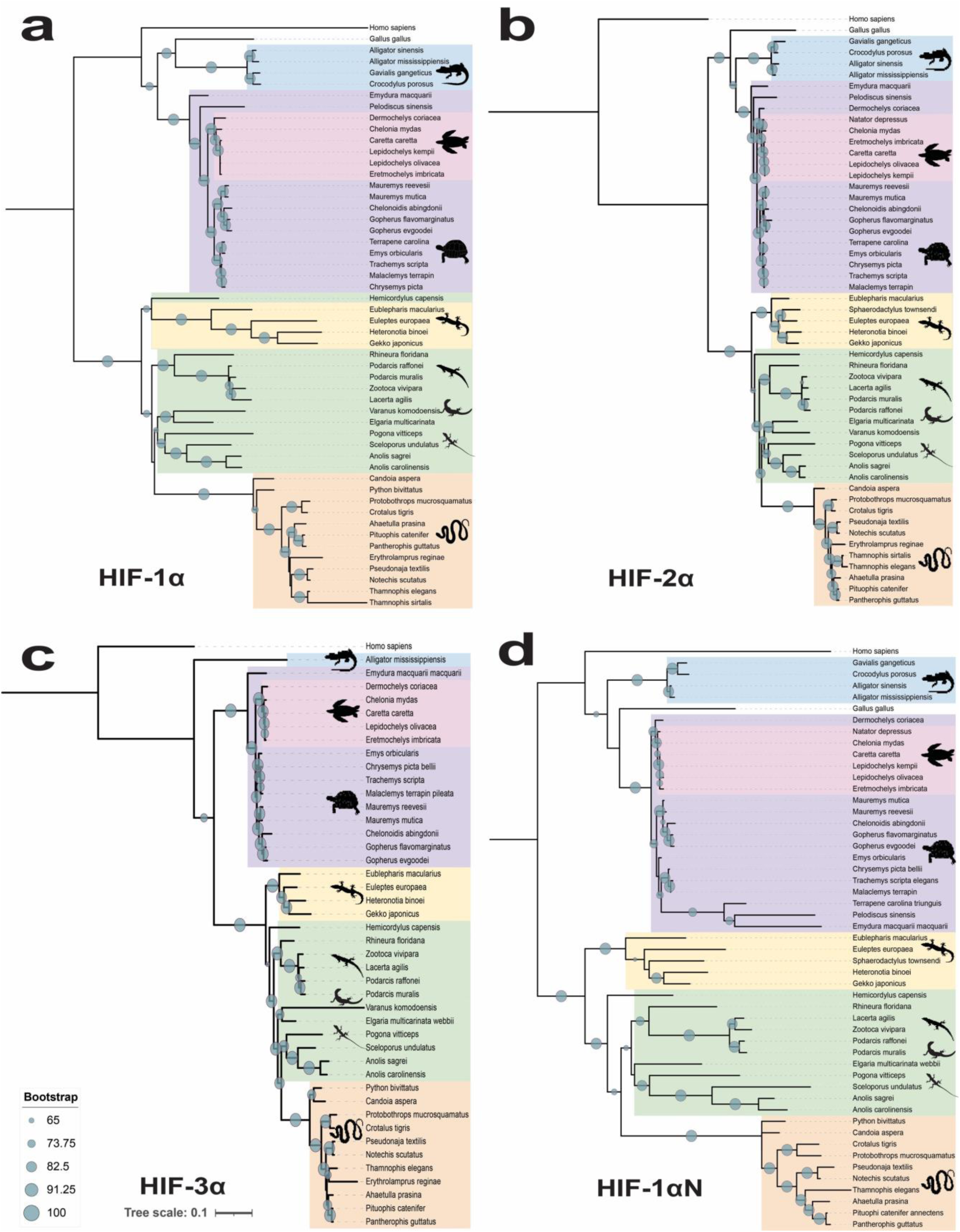
Phylogenetic reconstruction of HIF genes across reptiles. Maximum likelihood phylogenies of (a) HIF-1α, (b) HIF-2α, (c) HIF-3α, and (d) HIF-1αN across 36-50 reptile species. All trees show strong node support, and all genes seem conserved across species.

**Figure 3.**
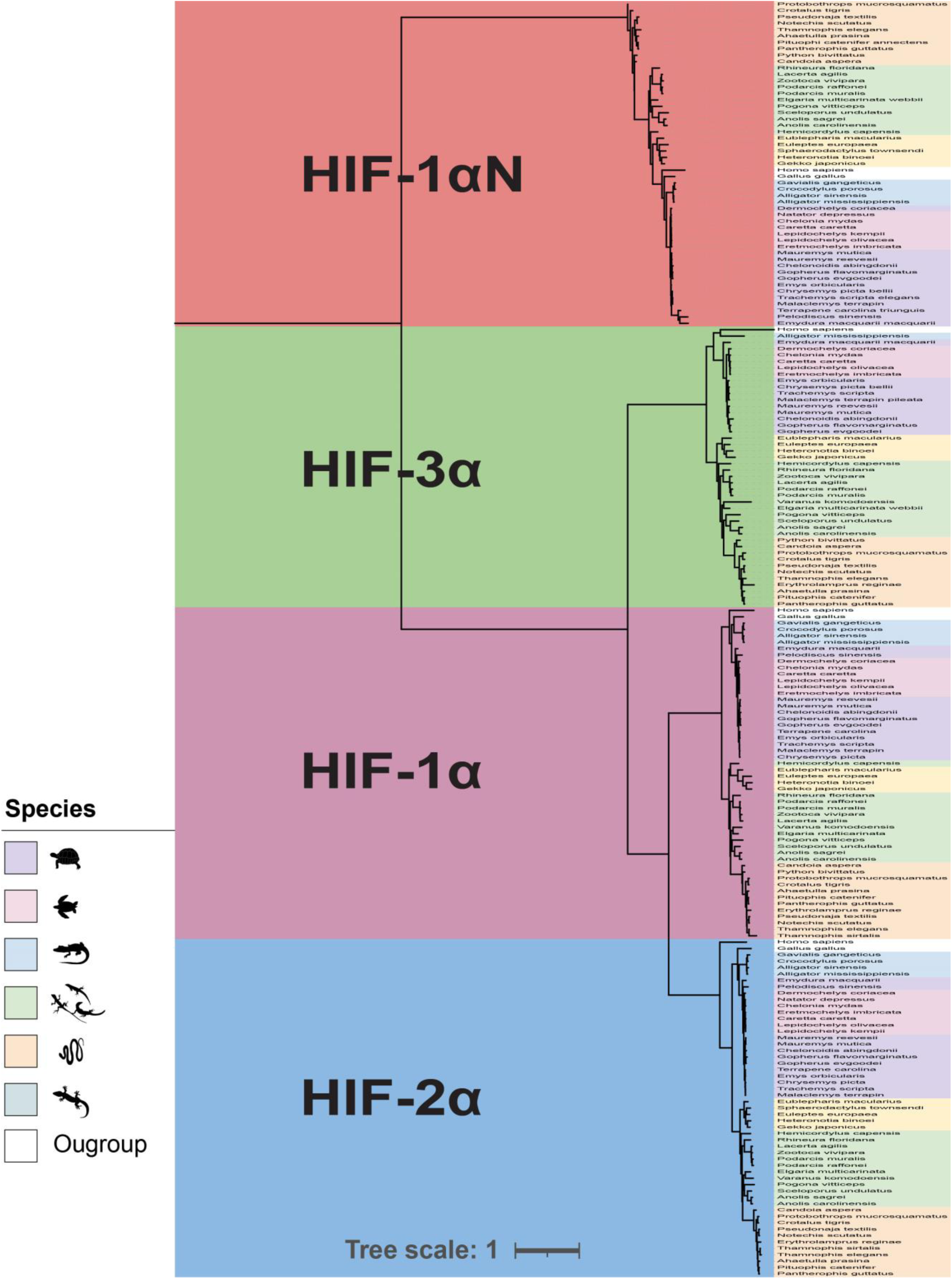
Evolutionary relationships among HIF genes. Combined phylogenetic reconstruction of HIF genes showing their relationship to each other. HIF-1αN and HIF-3α are most closely related, while HIF-2α represents the most recent divergence in the gene family.

We tested for positive selection on four HIF genes (HIF-1α, HIF-1αN, HIF-2α, and HIF-3α) using branch-site models in *codeml*, designating Testudines (turtles) as the foreground lineage and all other reptiles as the background. Likelihood ratio tests comparing the alternative model (ω > 1 allowed in Testudines) to the null model (ω fixed at 1) showed no evidence of positive selection of HIF-1α (LRT = 0, p = 1, kappa (ts/tv) = 3.36), HIF-2α (LRT = 0, p = 1, kappa (ts/tv) = 2.91), and HIF-1αN (LRT = 0, p = 1, kappa (ts/tv) = 2.33); **Table 2**). In contrast, HIF-3α showed a significant signal of positive selection in the turtle lineage (LRT = 11.21, p = 0.0008, kappa (ts/tv) = 4.11; **Table 2 and Supplementary Table 1**). However, no significant differences were found when testing whether sea turtle HIF genes are under a different selection regime relative to other reptiles (all LRT = 0, p = 1). These findings show that while HIF-1α, HIF-2α, and HIF-1αN are highly conserved across reptiles, positive selection of HIF-3α in Testudines might reflect adaptive evolutionary changes that support hypoxia tolerance in this Order.

**Table 2.** Branch-site model estimates of site class proportions and dN/dS (ω) values for HIF genes. The proportions of codon sites under purifying selection (class 0), neutral evolution (class 1), or potential positive selection, with Testudines (turtles) as the foreground lineage and all other reptiles as the background (classes 2a and 2b) are shown. Background and foreground ω values correspond to the dN/dS ratios estimated by codeml under model 2. Only HIF-3α showed statistically significant evidence of positive selection, with a foreground ω₂ > 1 in site classes 2a and 2b.

| Gene | Site Class | Proportion | Background $\omega$ | Foreground $\omega$ |
| --- | --- | --- | --- | --- |
| HIF-1α | 0 | 0.61440 | 0.06390 | 0.06390 |
|  | 1 | 0.12106 | 1.00000 | 1.00000 |
|  | 2a | 0.22100 | 0.06390 | 1.00000 |
|  | 2b | 0.04355 | 1.00000 | 1.00000 |
| HIF-2α | 0 | 0.61708 | 0.06473 | 0.06473 |
|  | 1 | 0.09911 | 1.00000 | 1.00000 |
|  | 2a | 0.24454 | 0.06473 | 1.00000 |
|  | 2b | 0.03927 | 1.00000 | 1.00000 |
| HIF-3α | 0 | 0.84063 | 0.05208 | 0.05208 |
|  | 1 | 0.08769 | 1.00000 | 1.00000 |
|  | 2a | 0.06491 | 0.05208 | **7.47101** |
|  | 2b | 0.00677 | 1.00000 | **7.47101** |
| HIF-1αN | 0 | 0.86416 | 0.02788 | 0.02788 |
|  | 1 | 0.11254 | 1.00000 | 1.00000 |
|  | 2a | 0.02061 | 0.02788 | 1.00000 |
|  | 2b | 0.00268 | 1.00000 | 1.00000 |

We identified five conserved functional motifs among reptilian HIF gene promoters, totaling 20. Testudines express four to five motifs per HIF gene, while other reptiles express none to three, suggesting novel transcription factor binding sites for HIF genes in Testudines (**Figure 4 and Supplementary Figures 3a-d**). In HIF-1α, three motifs matched known transcription factors in the C2H2 Zinc Finger and Fork Head/Winged Helix families. Two motifs in HIF-2α matched C2H2 Zinc Finger factors, while HIF-3α had three matched motifs—two in the C2H2 Zinc Finger family, and one in the SMAD/NF-1 DNA binding domain family. HIF-1αN had only one known motif match in the C2H2 Zinc Finger family. Hence, Testudines exhibited the highest number of motifs per HIF gene among reptiles, with conserved and lineage-specific patterns that may reflect the evolution of distinct transcriptional regulatory mechanisms for HIF genes in this lineage.

**Figure 4.**
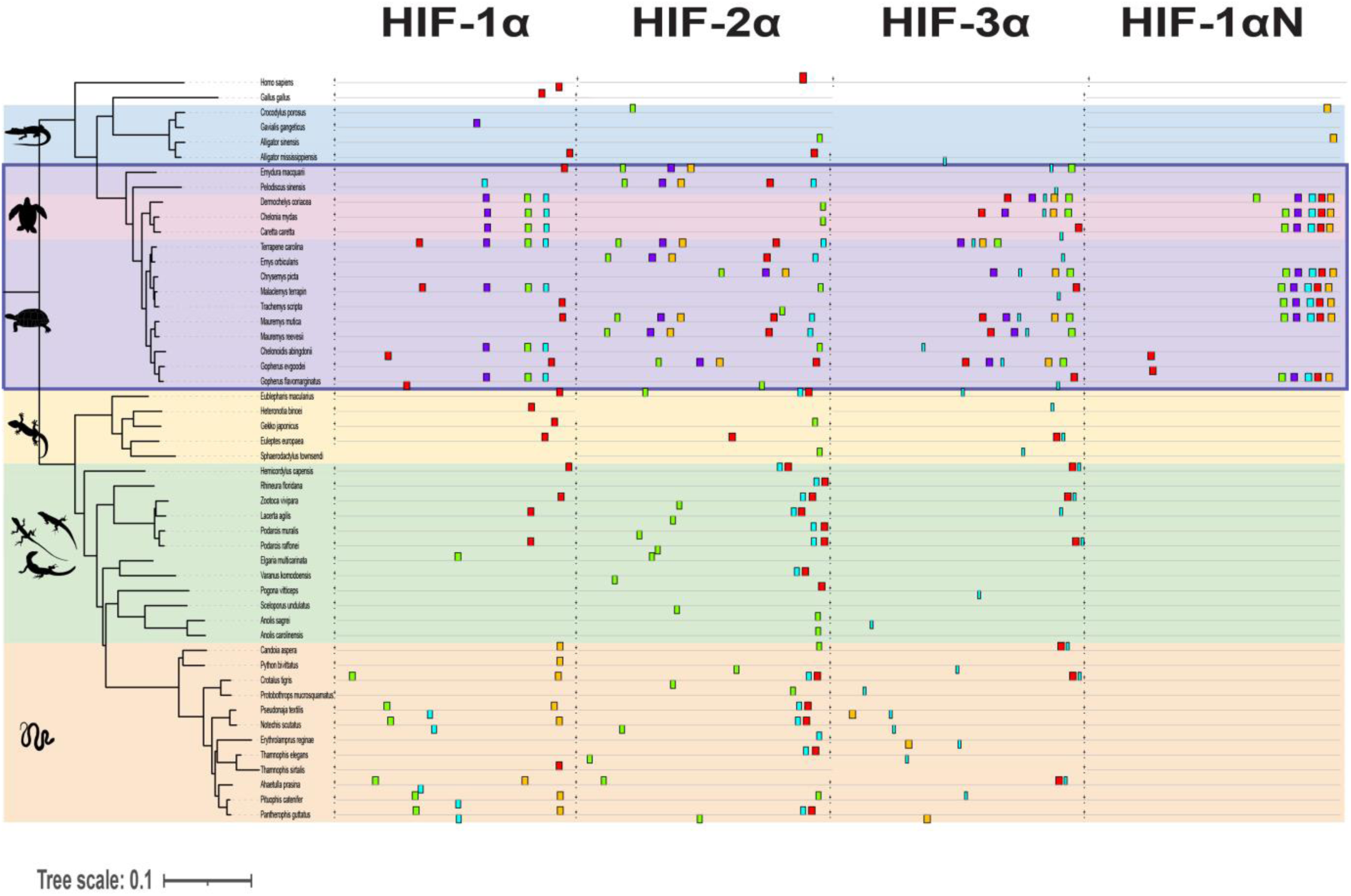
Promoter motif divergence among reptilian HIF genes. Testudines species exhibit a higher number of promoter motifs, 4-5 per gene, while other reptiles show only 1-3 motifs, suggesting the evolution of novel transcriptional regulatory elements in testudines.

### Gene expression trajectories follow species-specific patterns during short- and long-term hypoxia exposure in sea turtle and lizard cells

Besides inducing HIF transcription, which is crucial for the sustained response to hypoxia (Minet *et al*., 1999; BelAiba *et al*., 2007), low-oxygen exposure promotes HIF stabilization, increasing transcription of HIF-regulated genes that regulate oxygen transport and homeostasis, glucose metabolism, and cell survival (Wang *et al*., 1995; Semenza, 2008). Hence, we used RNA-seq to evaluate gene expression trajectories in sea turtle and lizard cells exposed to short-term (1-hour) hypoxia and reoxygenation. While we detected clusters of genes with similar expression trajectories in both species, we found distinctly enriched pathways. For instance, cluster 2 in lizard cells (**Figure 5a**) and cluster 1 in sea turtle cells (**Figure 5b**) both showed ascendant trajectories upon hypoxia exposure, followed by a return to baseline during reoxygenation; however, cluster 2 in lizard cells was enriched for pathways involved in cell repair, transcriptional repression, lipolysis, and DNA repair, whereas cluster 1 in sea turtle cells was enriched for pathways related to glycogen metabolism, mitochondrial translation, RNA degradation to manage oxidative stress (**Figure 6 and Supplementary Figure 4**). Similarly, cluster 5 in lizard cells (**Figure 5a**) and cluster 3 in sea turtle cells (**Figure 5b**) showed slightly downward trajectories during hypoxia exposure and rapid upregulation during reoxygenation. Here, in lizard cells, this cluster was enriched for developmental and cell function pathways, including cytoskeleton organization, chromatin remodeling, glycosaminoglycan biosynthesis, lysine degradation, and signaling pathways such as AGE-RAGE, Rho GTPases, and RORA-mediated transcription. In contrast, cluster 3 in sea turtle cells was enriched for pathways essential for cellular stress response and repair, including JAK-STAT, MAPK, neurotrophin signaling, transduction, TP53 regulation, mitochondrial quality control, and cellular homeostasis recovery (**Figure 6 and Supplementary Figure 4**). Thus, while we detected clusters of genes with similar gene expression trajectories during short-term hypoxia and reoxygenation in both species, genes clusters in lizard cells are enriched for pathways involved in rapid cellular stress responses. In contrast, in sea turtle cells, we detected enrichment for pathways targeting mitochondrial function and metabolic regulation, which likely help maintain homeostasis under hypoxic stress.

**Figure 5.**
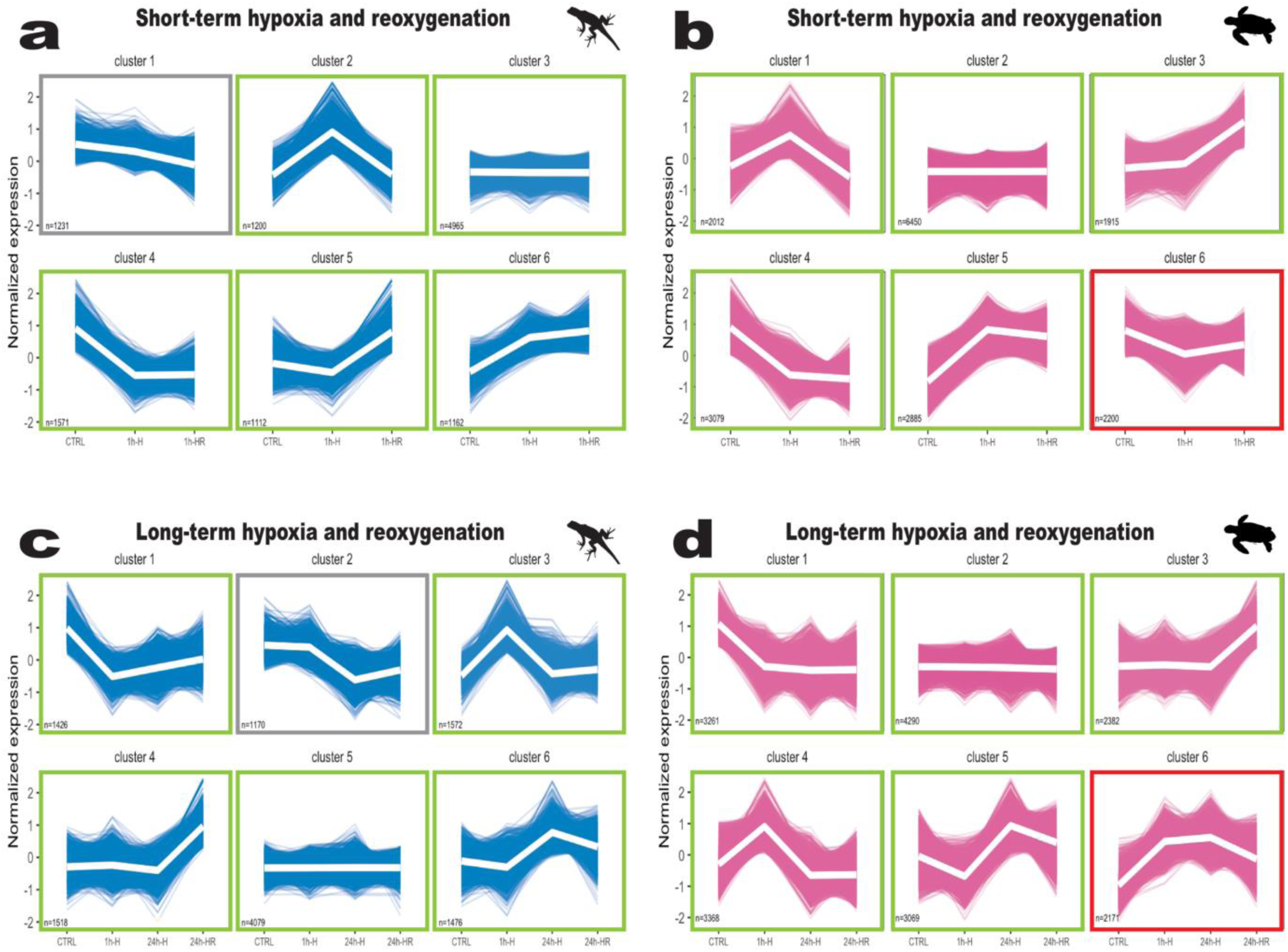
Gene expression trajectories in lizard and sea turtle cells exposed to hypoxia. Green borders indicate clusters with similar trajectories in both species. Red borders highlight unique trajectories in sea turtle cells. Clusters from control, 1h hypoxia, and 1h reoxygenation in (a) lizard cells and (b) sea turtle cells. Clusters from control, 1h hypoxia, 24h hypoxia, and 1h reoxygenation in (c) lizard cells and (d) sea turtle cells.

**Figure 6.**
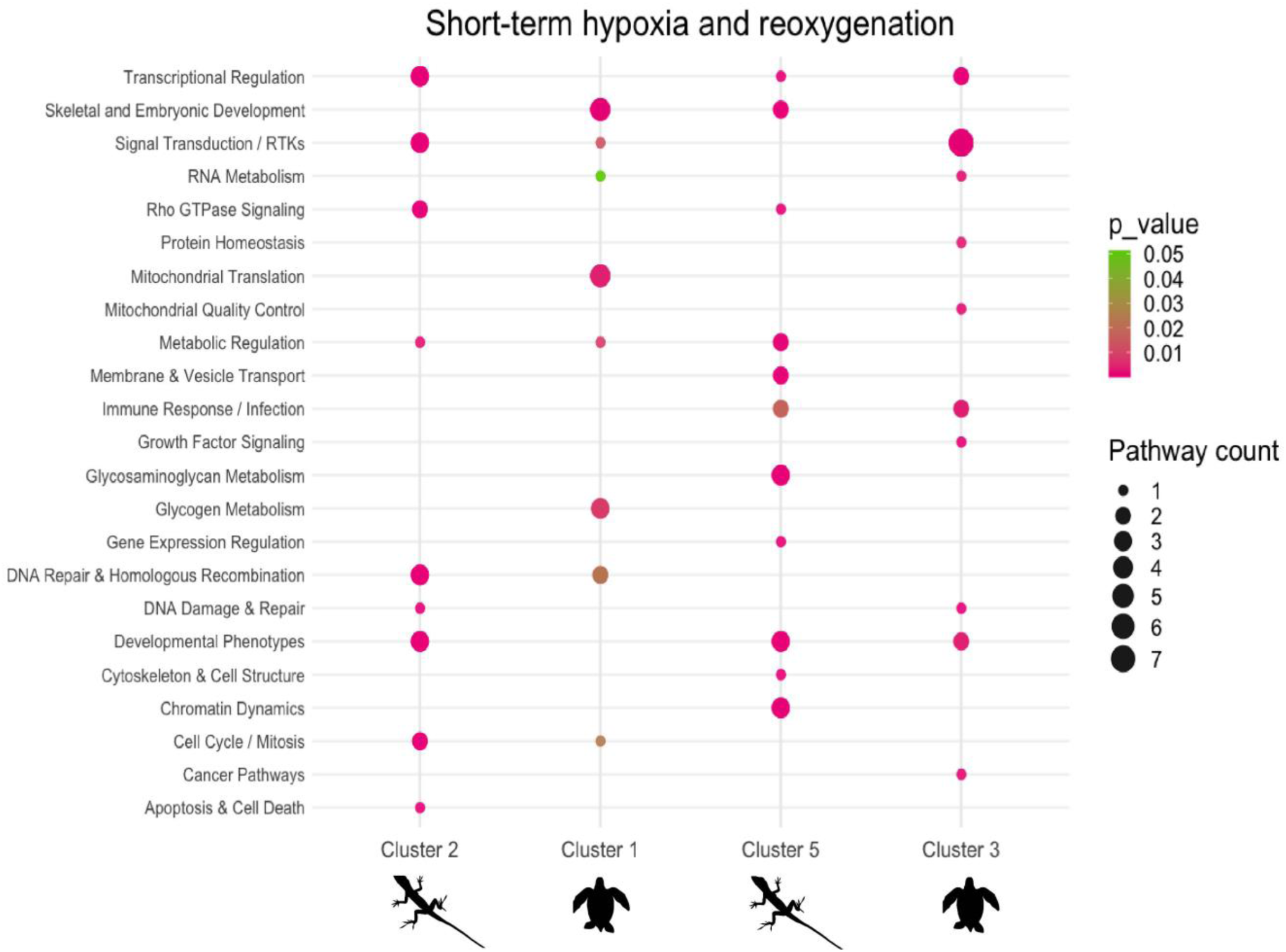
Functionally grouped enriched pathways from gene clusters (from. Figure 5**) that share similar expression trajectories during short-term hypoxia and reoxygenation in lizard and sea turtle cells**. Cluster 2 (lizard) mirrors the trajectory of Cluster 1 (sea turtle), while Cluster 5 (lizard) aligns with Cluster 3 (sea turtle).

We then examined the long-term (24-hour) response to hypoxia and reoxygenation to compare species-specific transcriptional profiles across timepoints. Both cluster 4 in lizard cells (**Figure 5c**) and cluster 3 in sea turtle cells (**Figure 5d**) showed no change from basal levels until reoxygenation, when expression trajectories increased. In lizard cells, enriched pathways include DNA repair, cell cycle regulation, and stress response mechanisms to DNA damage, replication stress, oxidative stress, and cell cycle disruption, which may help maintain genomic stability and coordinate cell growth and damage repair. In contrast, in sea turtle cells, we detected pathways enriched for protein synthesis, translation initiation, and membrane targeting, suggesting protein production and quality control to maintain cellular function (**Figure 7**). Hence, these results suggest that lizard cells respond to hypoxia and reoxygenation by prioritizing genes involved in energy homeostasis, DNA repair (homologous recombination and double-strand break repair), and cell survival. In contrast, sea turtle cells employ pathways involved in RNA processing, energy metabolism, mitochondrial function, and protein integrity. Of note, analyses of concerted changes in gene expression during both short- and long-term hypoxia/reoxygenation exposure revealed one cluster of genes with a markedly distinct trajectory in sea turtle cells that did not match any cluster in lizard cells (Cluster 6, **Figure 5b and 5d**). In this cluster, gene expression decreases at 1h and returns to basal levels with reoxygenation after 1h hypoxia. Enriched pathways in this cluster relate to RNA processing and transport—such as mRNA splicing via the spliceosome—involving genes including *EIF4A3, HNRNPR,* and *EFTUD2*, as well as protein synthesis and ribosome biogenesis processes, which might support cellular recovery by maintaining gene expression and proteostasis (**Figure 8a**). Sea turtle cells also show gene expression signatures related to preserving mitochondrial function, integrity, and redox homeostasis (*ISCA1, MTCH2, and RPTOR*), consistent with our previous work (Arango *et al*., 2025). Furthermore, the presence of *AKT1* and *RPTOR* suggests activation of mTOR-related pathways that enhance the activity of key antioxidant enzymes such as superoxide dismutase (SOD) and catalase (Glorieux *et al*., 2014; Tsang *et al*., 2018). During long-term hypoxia exposure, gene expression increases with hypoxia at 1-hour, stabilizes by 24-hours, and decreases upon reoxygenation. Enriched pathways in this cluster include Fox-O-mediated transcription involving FOXO3, PIK3R1, and IGF1R, which support antioxidant production and cellular repair (**Figure 8a-b**). Therefore, these unique transcriptional signatures enriched in sea turtle cells may bolster antioxidant defenses in preparation for reoxygenation, potentially mitigating oxidative damage, as observed in other animals adapted to cope with extreme oxygen fluctuations (Hermes-Lima *et al*., 2015; Arango *et al*., 2025).

**Figure 7.**
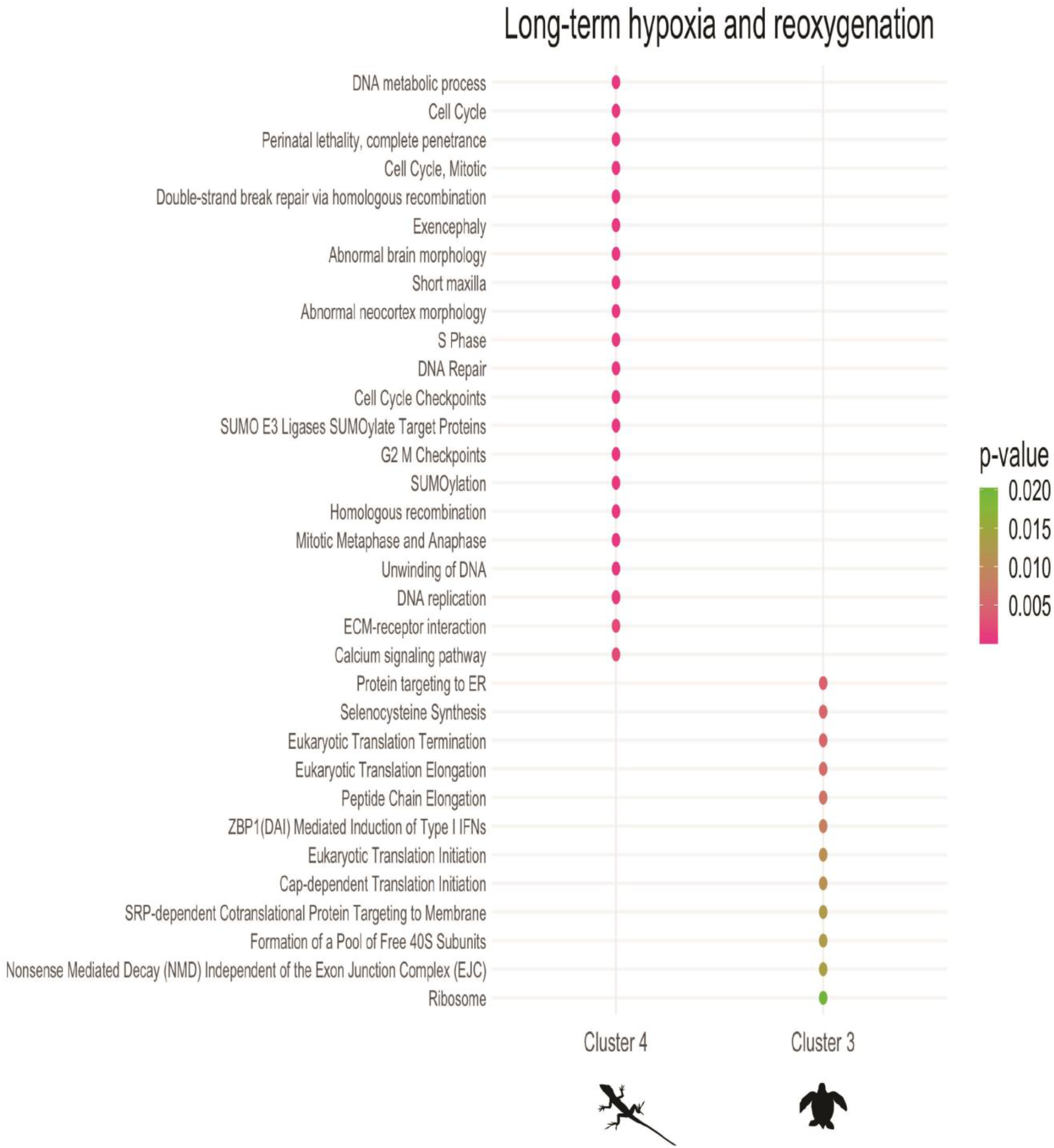
Enriched pathways in gene clusters with similar expression trajectories during long-term hypoxia and reoxygenation in lizard and sea turtle cells. Cluster 4 (lizard) aligns with Cluster 3 (sea turtle).

**Figure 8.**
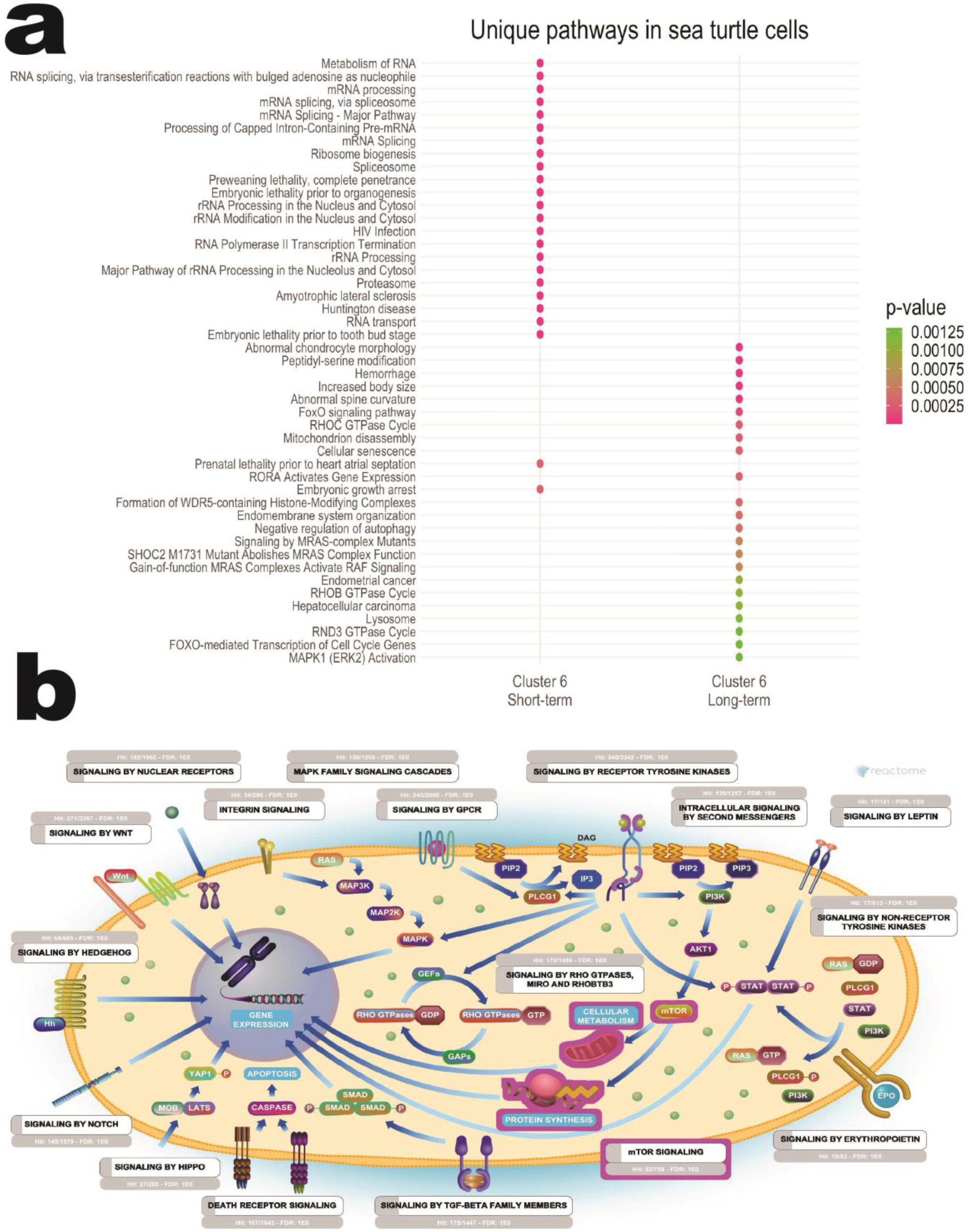
Enriched pathways from gene clusters with distinct expression trajectories during short- and long-term hypoxia and reoxygenation in sea turtle cells. (a) Cluster 6 from both short- and long-term treatments shows expression pathways not observed in lizard cells. (b) Reactome v93 FOXO-mediated MAPK pathway from cluster 6b.

### Sea turtle cells exhibit a robust transcriptional response to hypoxia compared to lizard cells

We next analyzed differentially expressed genes between control and each hypoxia/reoxygenation treatment in sea turtle and lizard cells. Overall, we detected thousands of DE genes in sea turtle cells. In contrast, we only detected tens to hundreds of DE genes in lizard cells. This order-of-magnitude difference suggests a more complex regulatory response to hypoxia in sea turtle cells than in lizard cells. The highest number of differentially expressed genes (1148 upregulated and 1415 downregulated) was observed in sea turtle cells exposed to 1-hour hypoxia/reoxygenation (**Figure 9**). This tightly regulated response to reoxygenation following short-term hypoxia exposure includes pathways enriched for ribosome biogenesis, rRNA and ncRNA processing, macromolecule synthesis, and metabolic processes, and functions related to protein synthesis, RNA processing, and cellular recovery, which are likely essential to restore homeostasis after hypoxic stress. In contrast, the fewest DE genes (34 upregulated and 18 downregulated) were detected in lizard cells after 24-hours of hypoxia exposure (**Figure 9**). Notably, no shared genes were upregulated between the two species during any hypoxia or reoxygenation time point, suggesting that reptilian transcriptional response to hypoxia is species-specific (**Figure 10**). However, 14 genes were downregulated in both species during short-term hypoxia and/or reoxygenation (**Figure 10**). Those shared genes are involved in metabolism (*FASN, TRIB3, MYC, NR4A3*), transcription and responses to oxidative, inflammatory and apoptotic stress (*CSRNP1, EGR2, ZC3H12A, TSC22D2, NCOR2*), cell structure and migration (*VIM, ZYX, CLIP1*), immune modulation (*ZC3H12A, NR4A3*), and extracellular matrix remodeling and signaling (*THBS2, ANKRD50*). These results suggest that downregulating these 14 shared genes may represent a minimal, conserved reptilian response to hypoxia, consistent with our cluster analysis, which showed similar gene expression trajectories but divergent enriched pathways between species. While lizard cells exhibit an attenuated transcriptional response to hypoxia, sea turtle cells regulate thousands of genes, suggesting a highly plastic response that may contribute to their natural tolerance to hypoxic stress.

**Figure 9.**
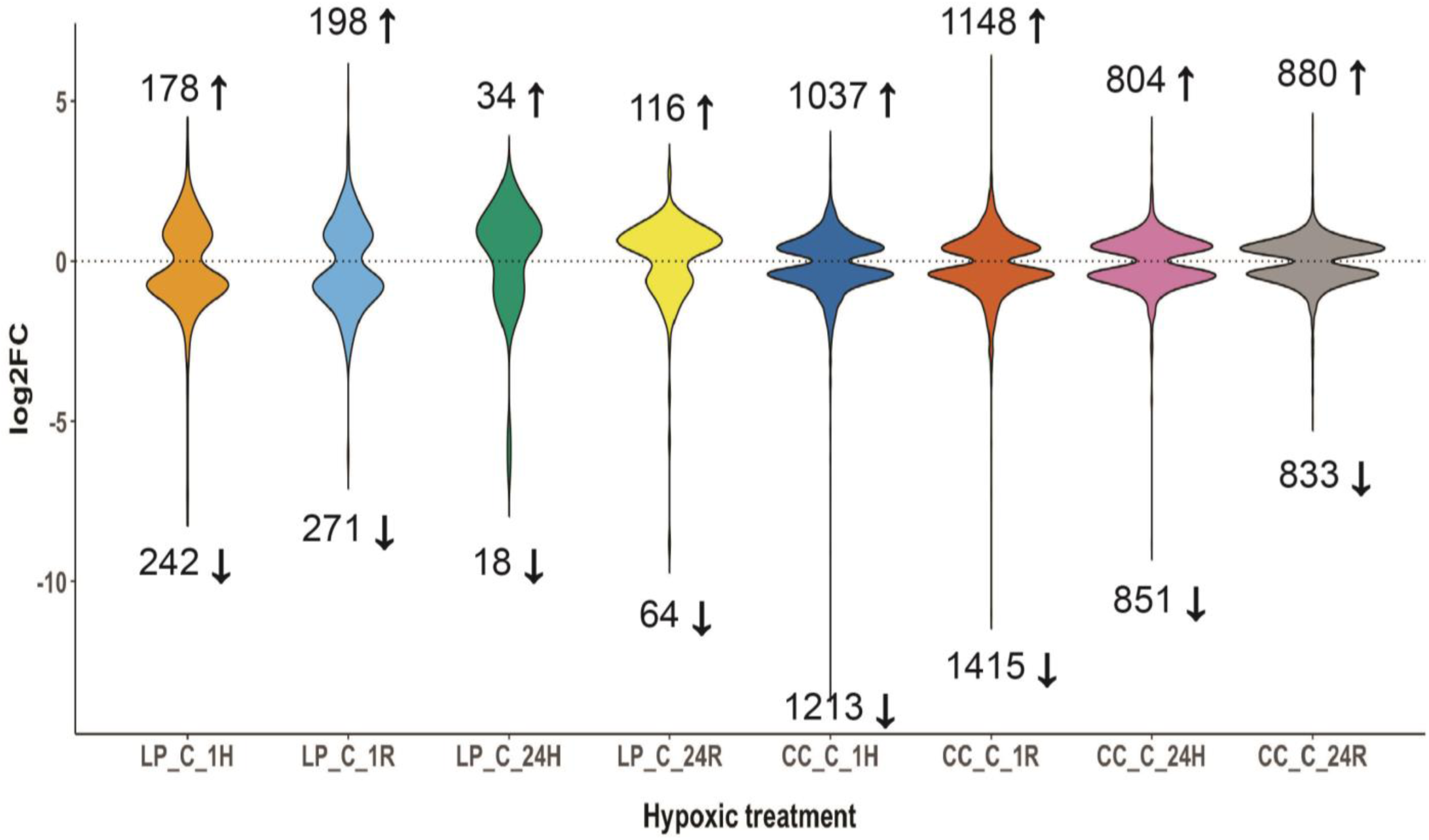
Differential gene expression between control and hypoxia/reoxygenation treatments. Sea turtle cells exhibit an order-of-magnitude increase in the number of differentially expressed genes compared to lizard cells, highlighting a markedly stronger transcriptional response to hypoxia. LP = lizard; CC = sea turtle; C = control; 1H = 1-hour hypoxia; 24H = 24-hour hypoxia; 1R = 1-hour reoxygenation following hypoxia; 24R = 24-hour reoxygenation following hypoxia.

**Figure 10.**
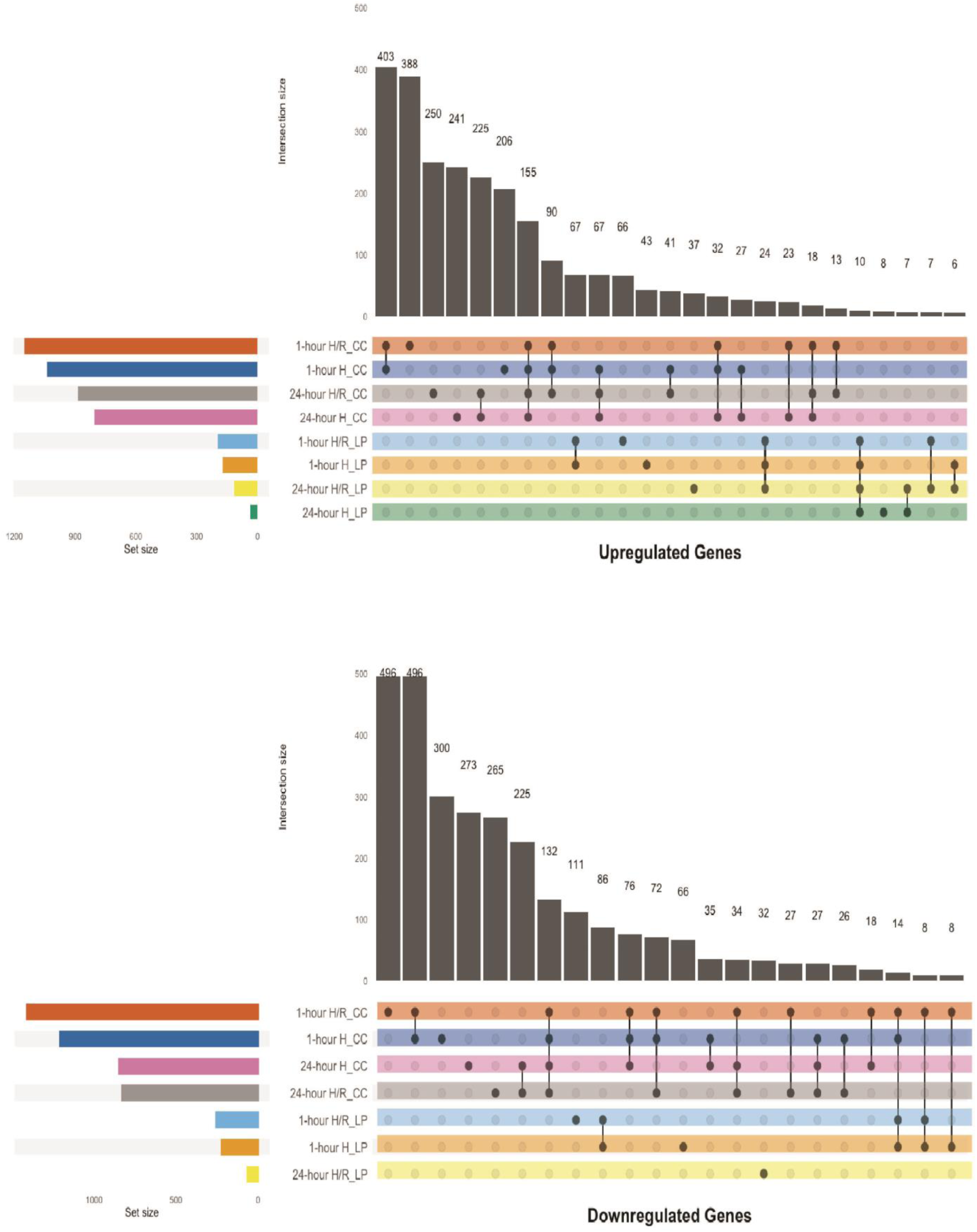
Differentially expressed gene interactions. UpSet plot summarizes the overlap of differentially expressed genes (DEGs) across hypoxia and reoxygenation treatments in lizard and sea turtle cells, highlighting species-specific upregulation and downregulation patterns. There are no shared genes upregulated in both species. There are 14 genes commonly downregulated in both species. Plot is showing > 5 gene interactions.

### HIF-1α might not be the primary transcriptional regulator of the cellular response to hypoxia in reptiles

Based on the full normalized gene expression dataset, the complete HIF gene family was detected in sea turtle cells, whereas only HIF-1α, HIF-2α, and HIF-3α were expressed in lizard cells. However, neither HIF-3α nor HIF-1αN were DE in either species, despite being known oxygen-dependent regulators (Zhang *et al*., 2014; Yang *et al*., 2015; Sim *et al*., 2018). HIF-1α was present among DE genes—in lizard cells, it is upregulated after 24 hours of hypoxia/reoxygenation, while in sea turtle cells, it is downregulated after 1-hour hypoxia/reoxygenation. In sea turtle cells, HIF-2α was upregulated after 24 hours of hypoxia/reoxygenation but downregulated after 1-hour.

We then mapped functional interaction networks (FIN) from each HIF transcription factor to our DE genes. FIN analysis identified only one known HIF-1α target gene (CXCR4) in lizard cells, compared to 34 targets in sea turtle cells, out of the hundreds of genes known to be regulated by HIF-1α (**Figure 11a**). Hence, these data suggest that although HIFs are typically considered the canonical master regulators of the transcriptional response to hypoxia (Guillemin & Krasnow, 1997; Semenza, 2008), this may not be the case in reptile cells.

**Figure 11.**
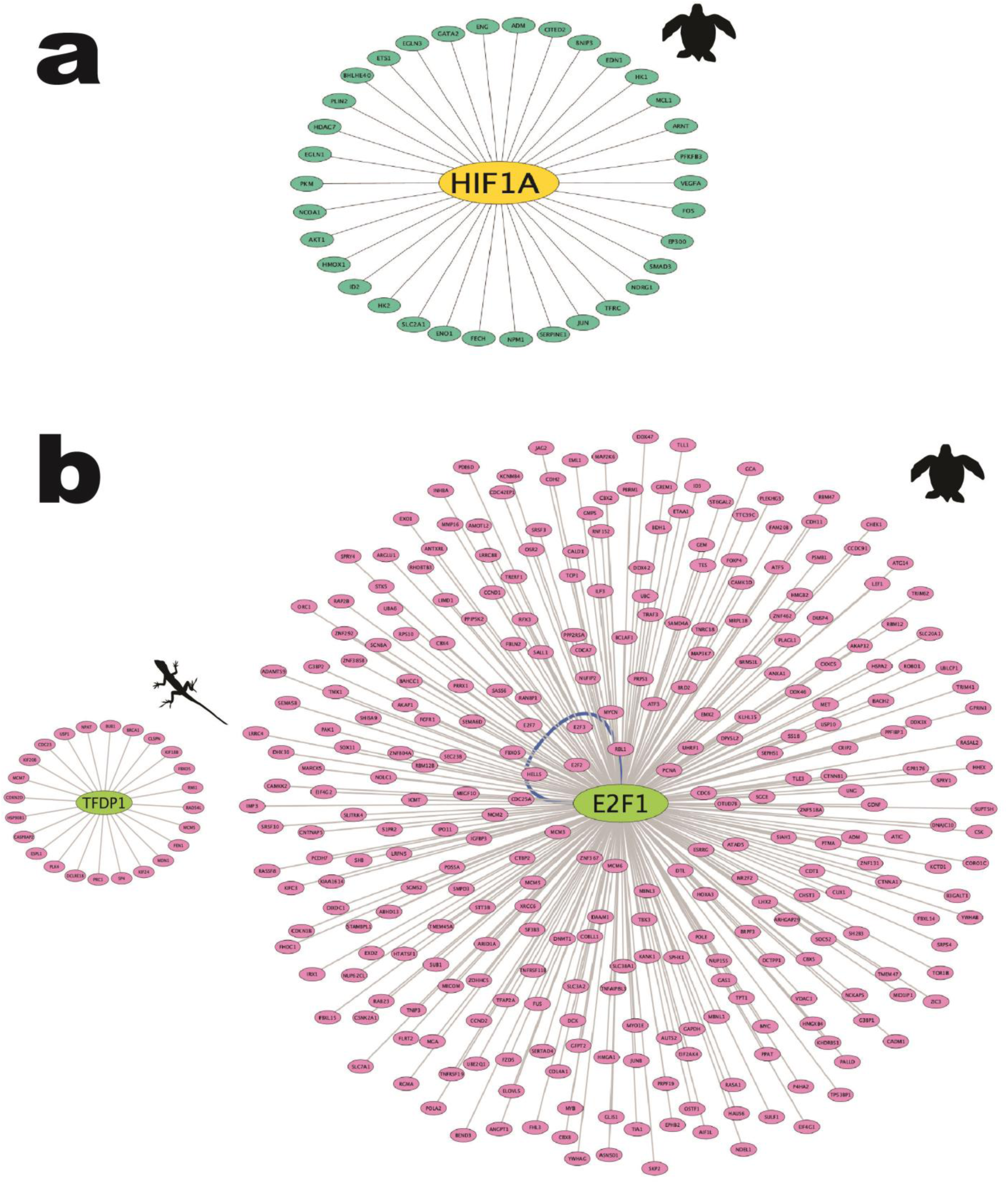
Functional interaction networks constructed with DE genes and transcription factors. (a) Sea turtle DE genes mapped against the known HIF-1α regulatory network. Only 34 genes are associated with known HIF-1α targets. (b) TFDP1 appears to be the main transcription factor driving DE of upregulated genes in the hypoxia response in lizard cells, with 24 genes mapping to its network. E2F1 appears to be the main transcription factor upregulating gene transcription in response to hypoxia in sea turtle cells, with 299 genes mapping to its network. The striking contrast in network size highlights the minimal transcriptional response in lizards compared to the extensive transcriptional regulation in sea turtle cells.

Next, we searched for *cis*-regulatory elements in all upregulated DE genes to identify potential transcription factors regulating these genes. In lizard cells, TFDP1 emerged as the most prominent transcriptional regulator, mapping to 24 genes with a normalized enrichment score (NES) of 6.45. This cluster of enriched motifs mapped exclusively to TFDP1. In contrast, E2F1 mapped to 299 genes in sea turtle cells (rather than 34 for HIF-1α) and had an NES of 5.87 (**Figure 11b**). This cluster included seven transcription factors identified by the motif2TF algorithm: E2F2, E2F3, E2F4, E2F5, E2F7, TFDP1, and TFDP3. Furthermore, mapping this entire cluster to all DE genes (downregulated and upregulated) accounted for 1179 associated genes. Thus, the primary transcription factors regulating DE genes in our dataset include members of the E2F and TFDP families, which are known to modulate cell cycle progression, DNA repair, and apoptosis—key processes for minimizing damage and ensuring cell survival under extreme oxygen fluctuations. Taken together, these results suggest that HIF-1α may not be the primary transcriptional regulator of the hypoxic response in reptile cells. Instead, TFDP1 in lizard cells and E2F1 in sea turtle cells appear to drive gene expression in hypoxic reptile cells. Moreover, these findings support the idea that lizard cells have a limited response to hypoxia, whereas sea turtle cells modulate gene expression to orchestrate a complex adaptive response.

## DISCUSSION

### HIF genes are mostly conserved across reptiles, but positive selection in HIF-3α and unique motifs identified in Testudines suggest adaptive transcriptional regulation

We examined whether HIF genes are conserved among reptiles or evolving under selective pressure. HIFs are key oxygen sensors that protect cells under low oxygen tension by activating metabolic regulators; therefore, their dysregulation can impair cell survival (Kumar & Choi, 2015). Similarly, under normoxia, HIF-1αN inhibits the transcriptional response to hypoxia. HIF-1αN also plays oxygen-independent functions in metabolic regulation (Majmundar *et al*., 2010; Zhang *et al*., 2011). Our phylogenetic analysis showed that HIF-1αN is the sister lineage to a clade containing HIF-3α, HIF-1α, and HIF-2α, while HIF-1α and HIF-2α share a more recent common ancestor with each other than to HIF-3α. Our results align with a previously reported vertebrate HIF phylogeny (Zhang *et al*., 2010), although that study excluded reptiles and HIF-1αN. Notably, we detected signatures of positive selection acting on HIF-3α in Testudines, which exhibit remarkable hypoxia tolerance (Hochscheid *et al*., 2005; Krivoruchko & Storey, 2015). This contrasts with previous studies in high-altitude human populations, where HIF-2α is under selection (Bigham *et al*., 2010; Simonson *et al*., 2010; Yi *et al*., 2010; Huerta-Sánchez *et al*., 2014; Jorgensen *et al*., 2023). Together, these findings suggest that while HIF-1α, HIF-2α, and HIF-1αN remain under strong purifying selection, likely due to functional constraints, HIF-3α may be adaptively evolving in Testudines in response to limited oxygen availability.

We next investigated whether alternative regulatory elements could initiate HIF transcription in Testudines. Transcription factor binding motifs serve as key regulatory elements that initiate gene expression (Bailey *et al*., 2015). We identified five promoter motifs in each HIF gene and found that Testudines possess more motifs compared to other reptiles. This suggests that testudines might have evolved distinct HIF transcriptional regulatory mechanisms unique from those of other reptiles. TomTom analysis grouped all HIF motifs into three major classes: C2H2 Zinc Finger Factors, Fork Head/Winged Helix Factor, and SMAD/NF-1 DNA Binding Domain Factors. The C2H2 Zinc Finger Factors family is the largest characterized transcription factor class, unique in its ability to bind long DNA sequences and interact with protein complexes, other transcription factors, and RNAs (Fedotova *et al*., 2017). Zinc-finger transcription factors (ZNFs) bind RNA and regulate post-translational processes (Nabeel-Shah *et al*., 2024). ZNFs influence gene expression and key processes, such as cell growth, differentiation, apoptosis, metabolism, and cellular stress responses (Mackeh *et al*., 2018). Among them, CTCF is a well-conserved transcriptional repressor critical for chromatin organization and imprinted gene regulation (Merkenschlager & Nora, 2016). Although CTCF is well conserved across bilaterians (Heger *et al*., 2012), CTCF appeared only in the HIF-1αN motif of Testudines, suggesting a lineage-specific role in hypoxia regulation. Given HIF-1α’s role as a master transcriptional regulator of hypoxia-responsive genes, these additional motifs may enable functional adaptations through transcriptional regulation of HIF-1α rather than coding sequence changes. Taken together, our motif analysis reveals novel transcription factor binding sites in testudines that may reflect a complex and coordinated adaptive transcriptional regulation of HIF genes in response to hypoxia.

### Distinct gene expression signatures underlie the cellular response to hypoxia in reptiles

Although three of four HIF genes are conserved across reptiles, motif analysis revealed potential novel transcriptional regulation of HIF gene in Testudines. Furthermore, our RNA data in hypoxic cells suggest distinct global regulatory strategies between sea turtle and lizard cells. The transcriptional response to hypoxia in turtle cells was strong, with thousands of differentially expressed genes (Gárriz *et al*., 2022; Wei *et al*., 2025). While gene expression pattern trajectories are broadly similar during short-term and long-term hypoxia in the two species, the underlying regulatory pathways appear species-specific. During short-term hypoxia and reoxygenation, gene expression signatures in lizard cells are enriched for pathways involved in rapid cellular stress response to hypoxia, whereas signatures in sea turtle cells are enriched for pathways related to mitochondrial function and metabolic regulation, which are likely to help maintain homeostasis under hypoxic stress. Under long-term hypoxia and reoxygenation, lizard cells upregulate genes involved in energy homeostasis, DNA repair, and cell survival. In contrast, DE genes in sea turtle cells are enriched for pathways related to RNA processing, energy metabolism, mitochondrial function, and protein integrity.

Notably, sea turtle cells uniquely activate Fox-O-mediated transcriptional pathways that promote antioxidant production and cellular repair, suggesting a preventative strategy to mitigate oxidative damage during reoxygenation (Reiterer & Milton, 2020; Jankovic *et al*., 2024). This mirrors findings in the Chinese yellow pond turtle (*Mauremys mutica)*, which survives 6-8 hours of hypoxia and exhibits DE genes enriched for inflammation, antioxidant defenses, P53, and MAPK signaling pathways under hypoxia (Wei *et al*., 2025). Similarly, in the hibernating Australian central bearded dragon (*Pogona vitticeps*), hypoxia and oxidative stress activate p53 and MAPK pathways (Capraro *et al*., 2019). Such strategies are also observed in other hypoxia-tolerant species, which preemptively upregulate antioxidant responses to reduce damage upon reoxygenation (Hermes-Lima *et al*., 2015; Arango *et al*., 2025). Prolonged hypoxia activates HIF-1α dependent transcriptional target genes, such as HIGD1A, that repress mitochondrial ROS production (Ameri *et al*., 2015). Consistent with this, we found that HIGD1A is downregulated in sea turtle cells during short-term hypoxia but upregulated after long-term hypoxia/reoxygenation. This pattern suggests low levels of mitochondrial ROS production, sufficient to initiate cell signaling, during short-term exposure, followed by repression during prolonged hypoxia to avoid oxidative damage. Thus, non-hypoxia-tolerant lizard cells likely mitigate oxidative damage through canonical responses that activate DNA repair and promote cell survival. In contrast, sea turtle cells, similar to other hypoxia-adapted vertebrates, likely adopt a preventative strategy by increasing antioxidant production and regulating mitochondrial architecture, function, metabolism, RNA integrity, and cellular homeostasis (Fabrizius *et al*., 2016; Reiterer & Milton, 2020; Allen *et al*., 2024). These findings are consistent with our previous work showing that sea turtle cells enhance antioxidant defenses and preserve mitochondrial integrity during hypoxia, which allows them to restart respiration during reoxygenation (Arango *et al*., 2025). These findings align with Jain et al., 2020, who demonstrated that regulating mitochondrial and lipid metabolism is essential for mounting an effective hypoxic response.

### Are HIFs the primary transcriptional regulators in hypoxic reptile cells?

Sea turtle cells mount a robust transcriptional response to hypoxia, regulating thousands of genes, whereas lizard cells show minimal regulation, highlighting a markedly more complex regulatory strategy in sea turtles. The highest number of DE genes in sea turtle cells occurs during reoxygenation after 1-hour hypoxia exposure, during which pathways related to ribosome biogenesis, rRNA, ncRNA processing, macromolecule synthesis, metabolic activity, protein synthesis, RNA processing, and cell recovery, all of which serve to restore homeostasis, are enriched. Hypoxia triggers damaging protein modifications, prompting cells to adapt through HIF-mediated transcription, mitochondrial regulation, and metabolic reprogramming (Lee *et al*., 2020). Lizards respond by repressing protein synthesis to mitigate oxidative damage. In contrast, sea turtle cells maintain protein folding and quality control under hypoxia, an active strategy that may help prevent oxidative damage, yet support continued function. Interestingly, there are only 14 downregulated genes commonly shared among the two species, most of which are associated with metabolism, transcriptional and cellular stress responses, cell structure and migration, immune modulation, extracellular matrix remodeling, and signaling. These may represent the minimal conserved hypoxia response in reptiles. Thus, while lizard cells exhibit a limited response, sea turtle cells regulate thousands of genes, suggesting a highly plastic and adaptive transcriptional response. However, given the limited taxonomic sampling, it remains unclear whether this represents a derived adaptation in turtles or in a larger reptilian clade. Broader comparative sampling will be essential to clarify whether the observed turtle response reflects lineage-specific hypoxia tolerance or a more ancestral reptilian response.

We found no evidence of HIF-3α or HIF-1αN DE in either species (downregulated or upregulated). Since HIF-1α functions predominantly as a transcriptional activator, any repression under hypoxia is likely mediated by HIF-1α-dependent induced repressor proteins and/or non-coding RNAs (Yfantis *et al*., 2023). This might partially explain our HIF-1α results for DE genes, where HIF-1α was upregulated after 24-hours hypoxia/reoxygenation in lizard cells but downregulated in sea turtle cells after 1-hour hypoxia/reoxygenation. We found limited HIF-1α-driven DE genes in our RNA-seq data, with only one HIF-1α target (CXCR4) in lizard cells and 34 in sea turtle cells, based on mapping between the HIF-1α network and all DE genes. These findings contrast with data from green sea turtles (*Chelonia mydas*) embryos during hypoxia induced by ovipositional arrest, where 57 and 27 DE genes mapped to known HIF-1α and HIF-2α targets (Gárriz *et al*., 2022). However, this discrepancy may be due to differences in hypoxia exposure, whereas our maximal exposure time was 24 hours, their study time point was 36 hours. This suggests that canonical HIF pathways in sea turtle cells might only be activated after extremely prolonged hypoxia exposure. Supporting this, ovipositional arrest in olive ridley sea turtles (*Lepidochelys olivacea*) can last 8-15 days (Williamson *et al*., 2019).

In contrast to our results, in *C. mydas* embryos exposed to 36-h hypoxia, both the HIF coactivator p300/CBP, and HIF-3α were upregulated (Capraro *et al*., 2019). HIF-1α has two transactivation domains (N-TAD and C-TAD); the C-TAD interacts with co-activators CBP/p300 to modulate gene transcription under hypoxia. Yet in our study, genes DE in sea turtles mapped only to p300, not CBP. Notably, DE of either CBP or p300 was not detected in lizard cells. Although CBP/p300 are functionally similar (Zeng *et al*., 2008), they have distinct functions, p300 acetylates HIF-1α, stabilizing the protein for a more sustainable HIF-1 transactivation (Geng *et al*., 2012). This may explain why hypoxic sea turtle cells show rapid nuclear accumulation of HIF-1α, unlike lizard cells (Arango *et al*., 2025).

Our transcription factor analysis identified TFDP1 as the primary transcriptional regulator of DE genes in hypoxic lizard cells, mapping to 19 genes. In contrast, E2F1 mapped to 299 genes in sea turtle cells. This module includes seven transcription factors from the E2F family. Members of the E2F and TFDP families modulate cell cycle progression, DNA repair, and apoptosis—key processes for minimizing damage and ensuring cell survival under oxygen fluctuations (Huang *et al*., 2021; Nakajima *et al*., 2023). TPD-1 and E2F1 form a crucial complex in regulating cell cycle control. These results suggest that HIF-1α may not be the primary transcriptional regulator of the hypoxic response in reptile cells. Instead, TFDP1 appears central in lizards, and E2F1 in sea turtle cells. Furthermore, the contrasting roles of these transcription factors support our conclusions that sea turtle cells manage hypoxia efficiently since antioxidant, mitochondrial, and metabolic regulation likely contribute to the efficient restoration of cell function during reoxygenation. Altogether, these findings suggest that TFDP1 and E2F1, in addition to HIFs, play a central role regulating the transcriptional response to hypoxia in reptile cells.

## CONCLUSSIONS

Our findings show that while HIF genes are mostly conserved among reptiles, positive selection on HIF-3α in Testudines may reflect adaptations to hypoxia. The transcriptional response to hypoxia is species-specific in reptile cells. Whereas lizard cells exhibit a canonical response focused on cell-survival, sea turtle cells respond to hypoxia by inducing a robust transcriptional program that regulates protein synthesis, quality maintenance, and mitochondrial integrity. These differences appear to be driven more by regulatory mechanisms than genetic divergence, underscoring the important role of transcriptional plasticity in reptilian hypoxic adaptation. Notably, HIF seems to play a limited role, with TFDP1 and E2F1 emerging as potential key regulators of the hypoxic response in reptiles.

## ACKNOWLEDGMENTS

We thank Noemi Bautista, Johnny Hoang, Yvette Javier, Mohamed Moustafa, Joseph Nguyen, Amber Singh, and Harshmeet Singh for their help with lizard collection and husbandry. Federico Kong-Gonzalez, Eqlima Tahiry and Dua Shoaib helped with wet lab analyses. Thank you to Dr. Jimmy McGuire for your help with phylogenetic discussions. We are also grateful to Dr. George Brooks for providing lab space access. BGA was supported by Ford Foundation predoctoral and dissertation fellowships. JPV-M is supported by NIGMS grant R35GM146951. Research funded by UC Berkeley.

## SUPPLEMENTARY MATERIALS

**Supplementary Table 1.** Positively selected sites in HIF-3α. Positively selected sites identified by Bayes Empirical Bayes (BEB) analysis (Yang *et al*., 2005). Only codon positions with BEB posterior probability greater than 0.95 (Prob(w > 1) > 0.95) are included.

| Position | Amino Acid | BEB Probability |
| --- | --- | --- |
| 22 | Q | 0.976 |
| 138 | E | 0.967 |
| 298 | G | 0.970 |
| 304 | E | 0.979 |
| 306 | N | 0.952 |
| 307 | P | 0.986 |
| 357 | R | 0.963 |
| 508 | D | 0.959 |
| 515 | P | 0.981 |
| 516 | P | 0.984 |
| 517 | T | 0.985 |

**Supplementary Figure 1.**
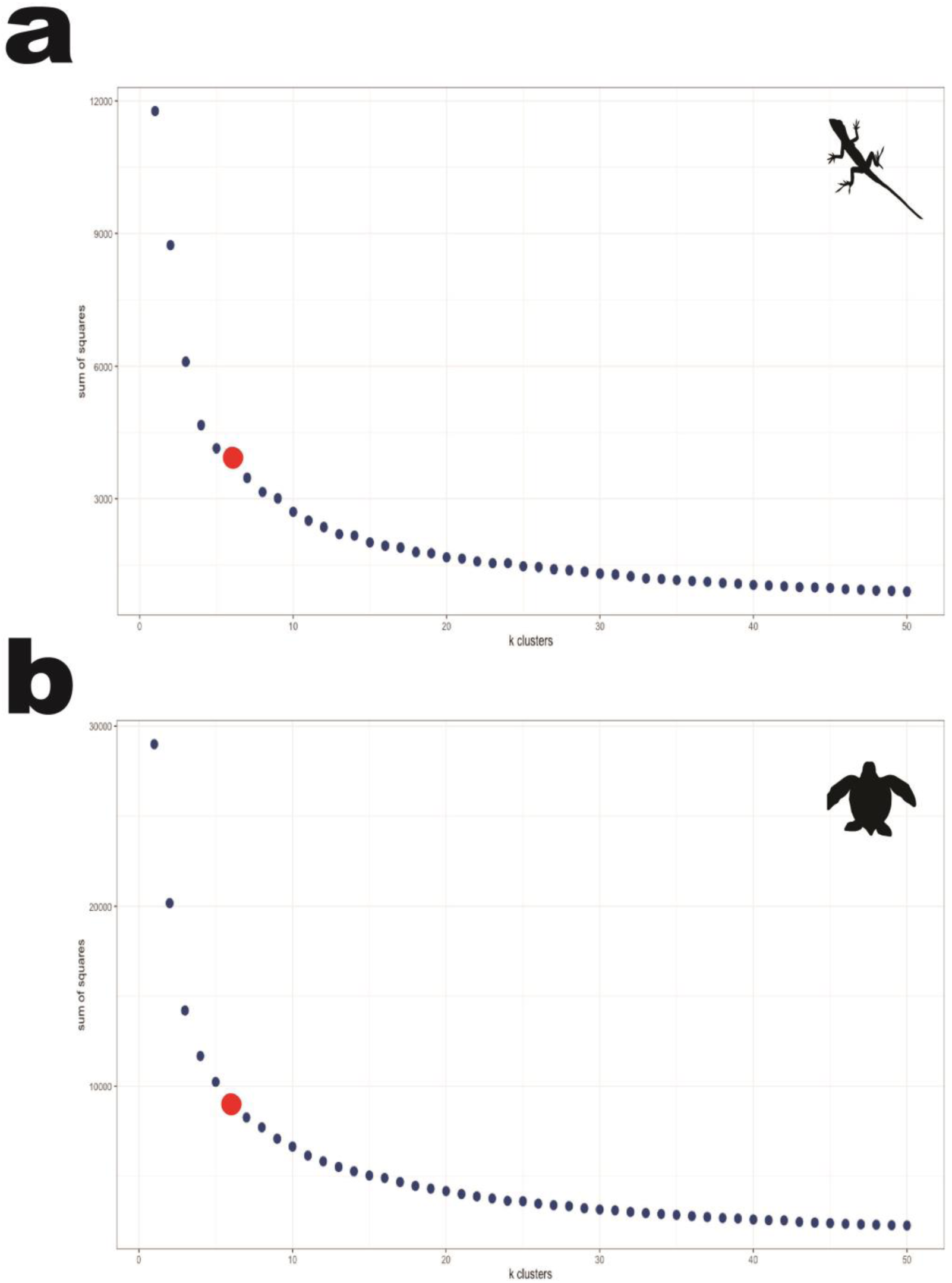
Elbow plots using silhouette scores to select the optimal number of clusters. (a) Lizard cells and (b) sea turtle cells: optimal cluster number set at the 6th inflection point from scores across 1-50 iterations.

**Supplementary Figure 2.**
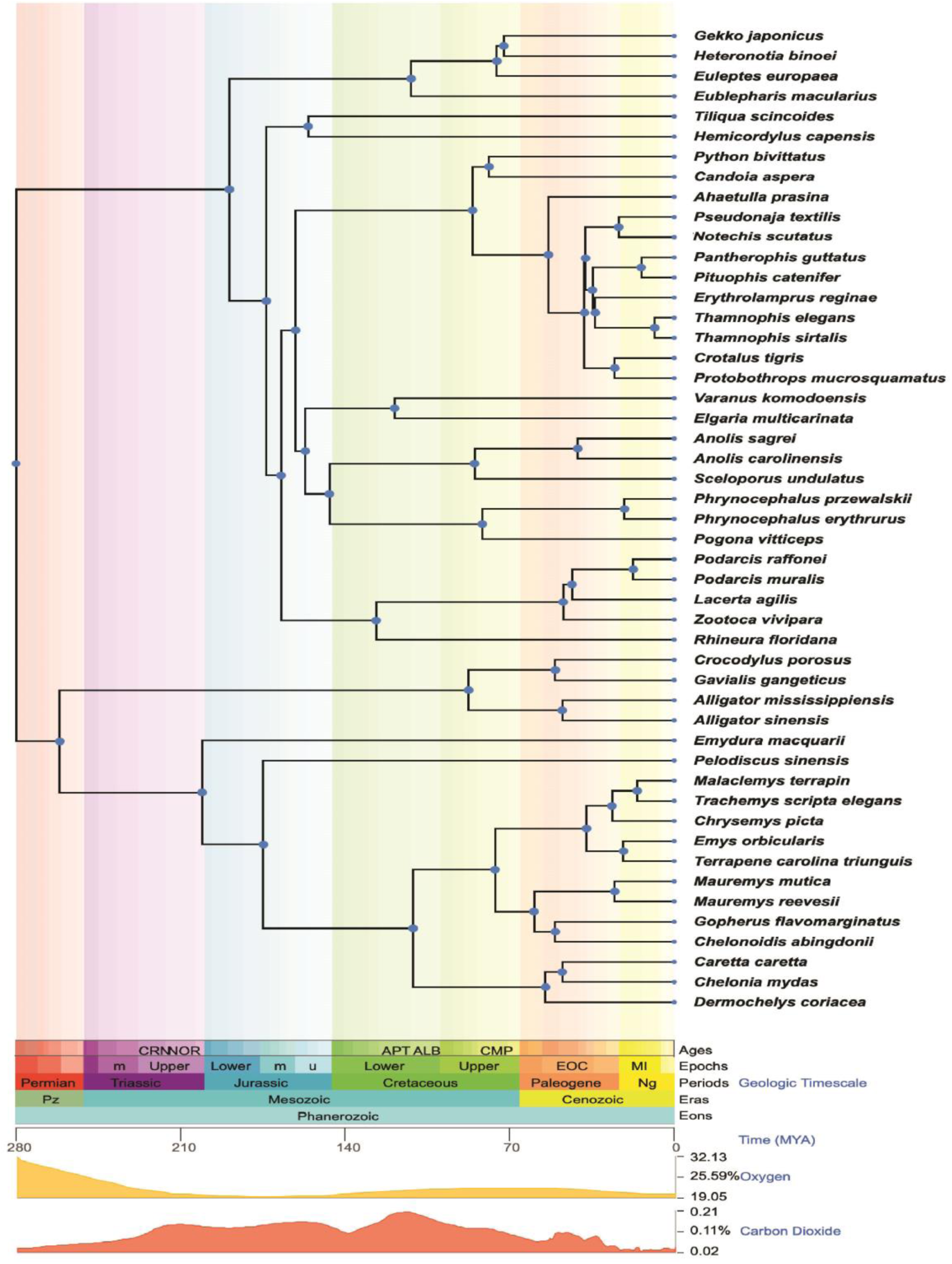
Testudines ancestors evolved during the late Permian. Time-calibrated reptilian species tree in TimeTree (https://timetree.org/) mapped to the geological time scale, major Earth impact events, and fluctuations in atmospheric oxygen and carbon dioxide levels.

**Supplementary Figure 3a.**
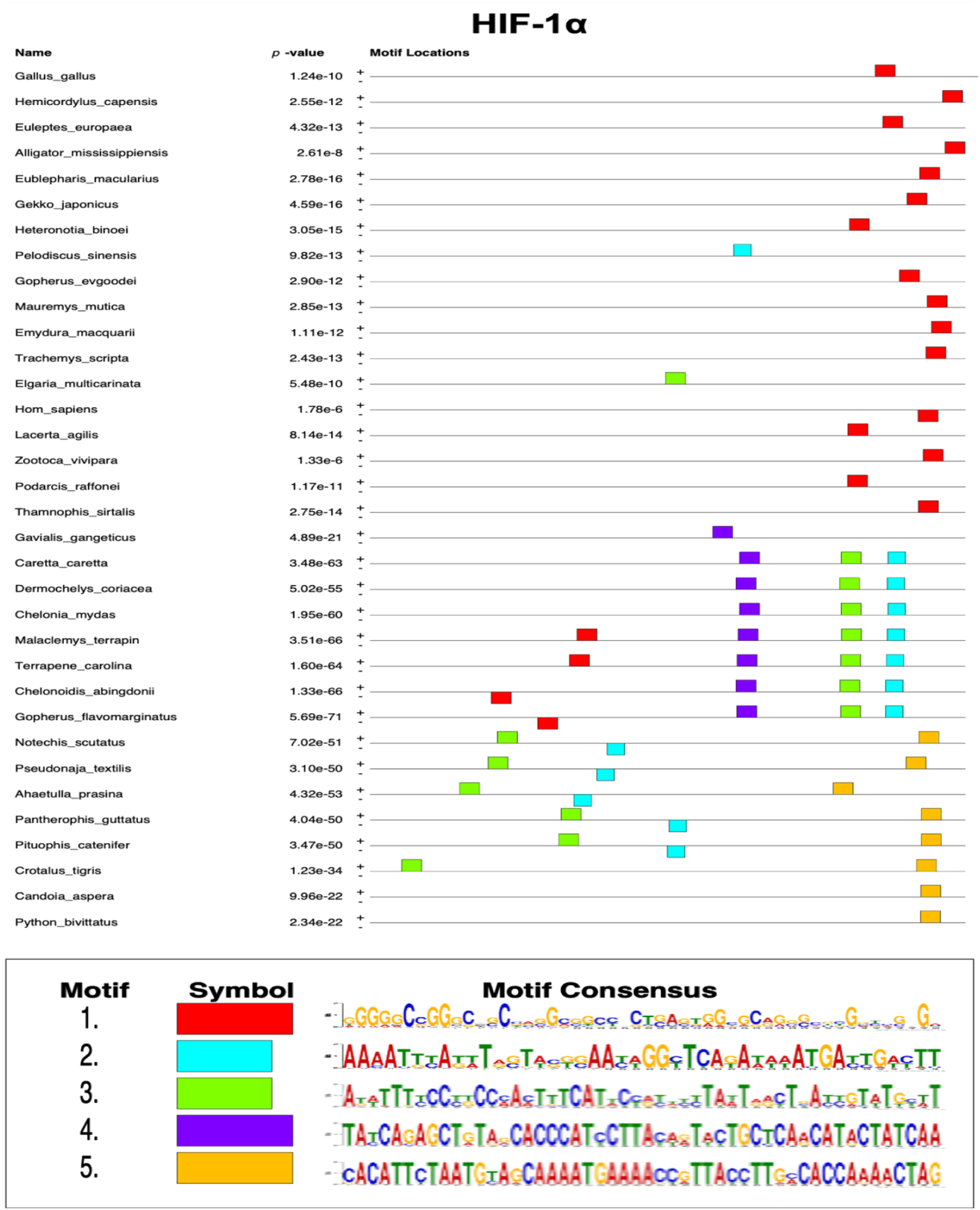
Predicted transcription factor binding motifs in HIF-1α promoter regions across reptiles. Motifs are shown with their respective E-values, positional enrichment, and sequence logos. Testudines (turtles) exhibit a greater number and diversity of conserved motifs compared to other reptilian lineages.

**Supplementary Figure 3b.**
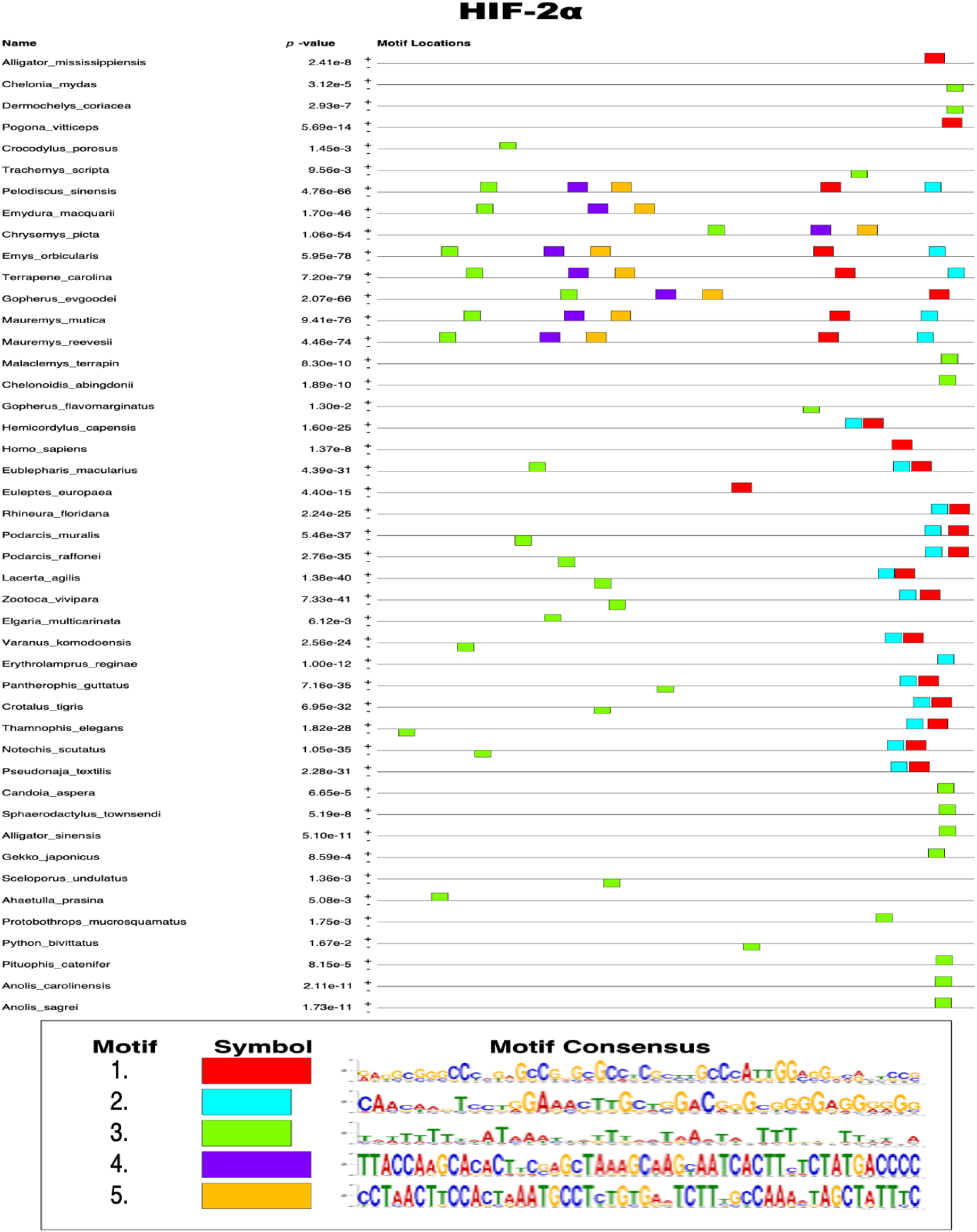
Predicted transcription factor binding motifs in HIF-2α promoter regions across reptiles. Motifs are shown with their respective E-values, positional enrichment, and sequence logos. Testudines (turtles) exhibit a greater number and diversity of conserved motifs compared to other reptilian lineages.

**Supplementary Figure 3c.**
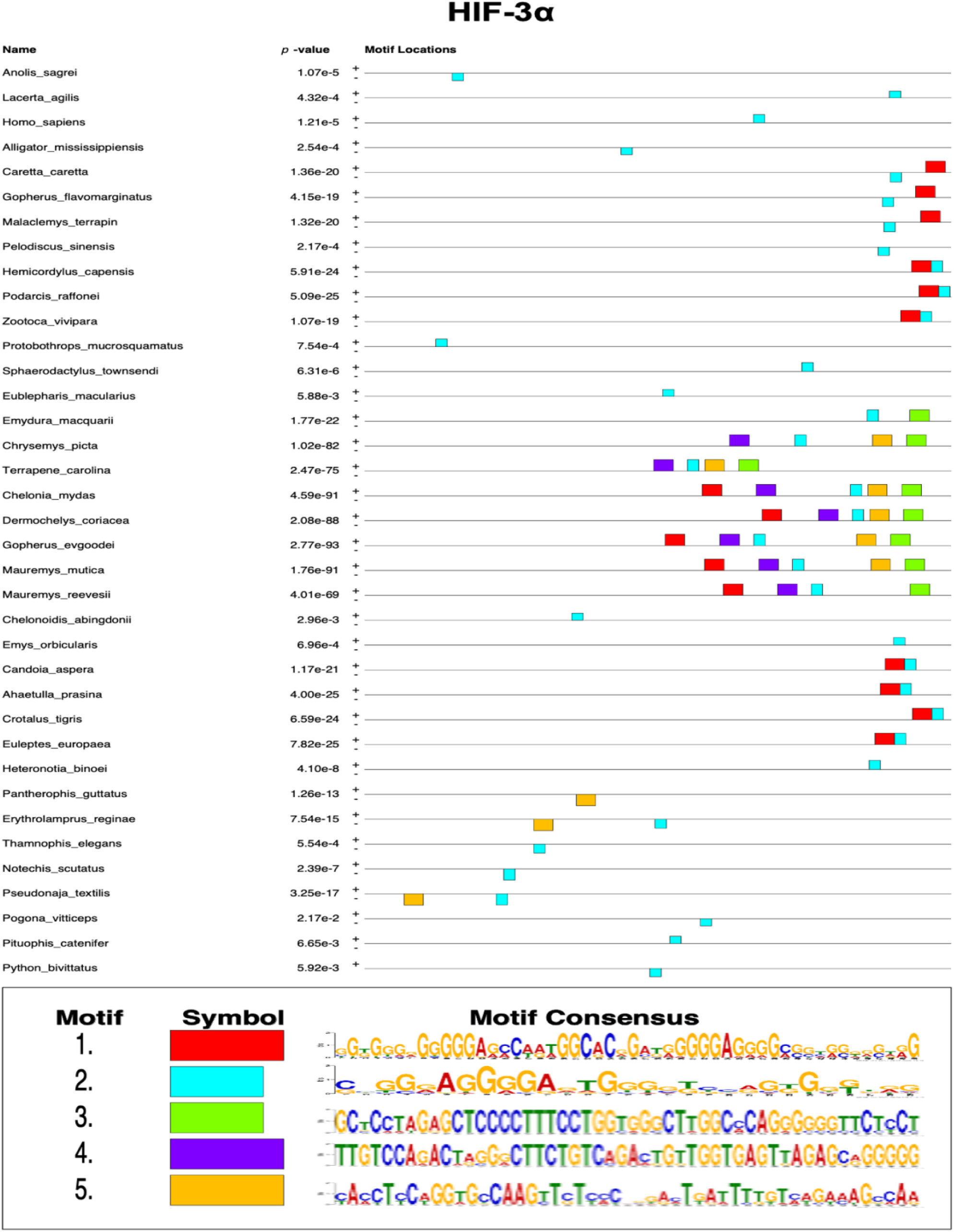
Predicted transcription factor binding motifs in HIF-3α promoter regions across reptiles. Motifs are shown with their respective E-values, positional enrichment, and sequence logos. Testudines (turtles) exhibit a greater number and diversity of conserved motifs compared to other reptilian lineages.

**Supplementary Figure 3d.**
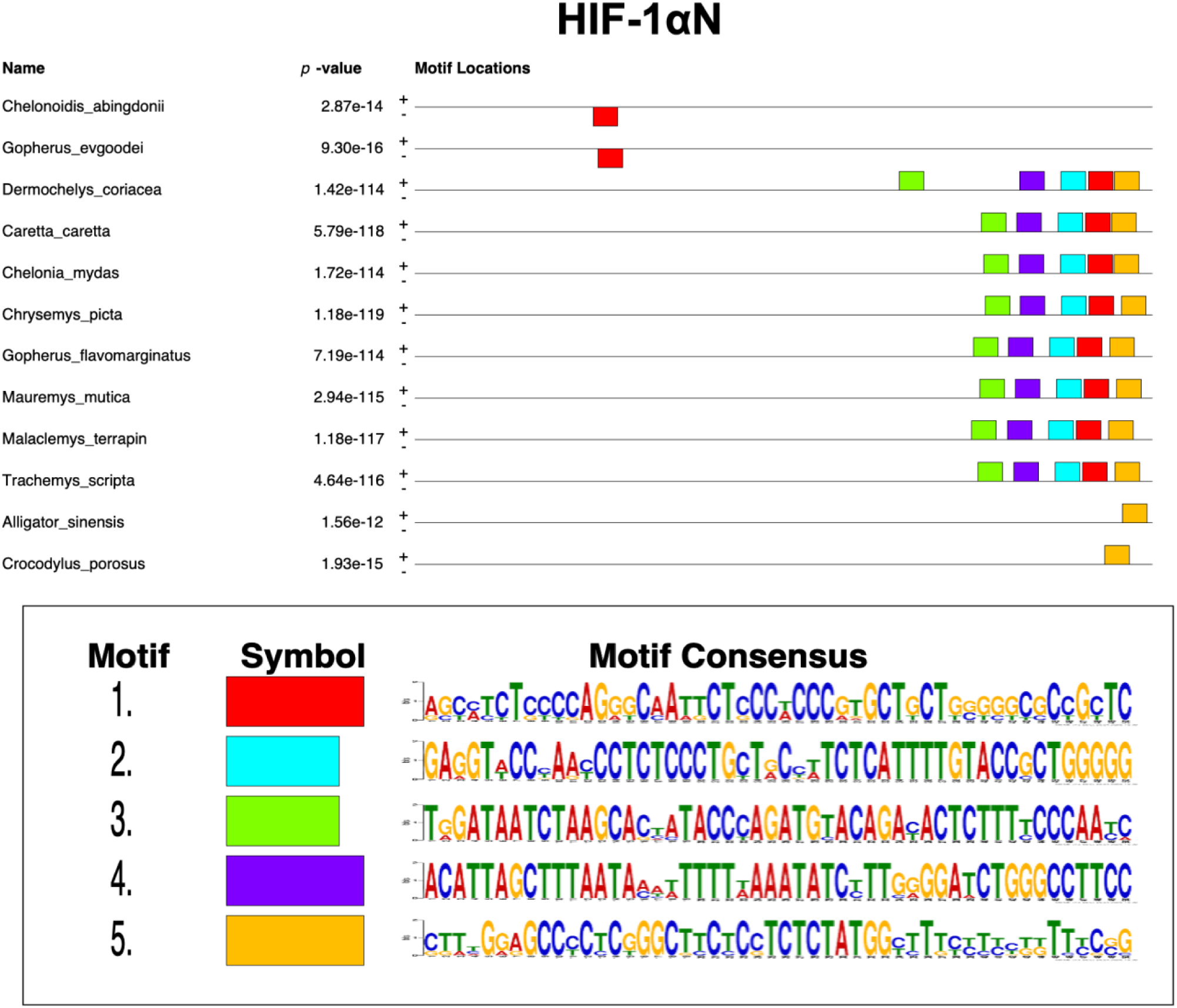
Predicted transcription factor binding motifs in HIF-1αN promoter regions across reptiles. Motifs are shown with their respective E-values, positional enrichment, and sequence logos. Testudines (turtles) exhibit a greater number and diversity of conserved motifs compared to other reptilian lineages.

**Supplementary Figure 4.**
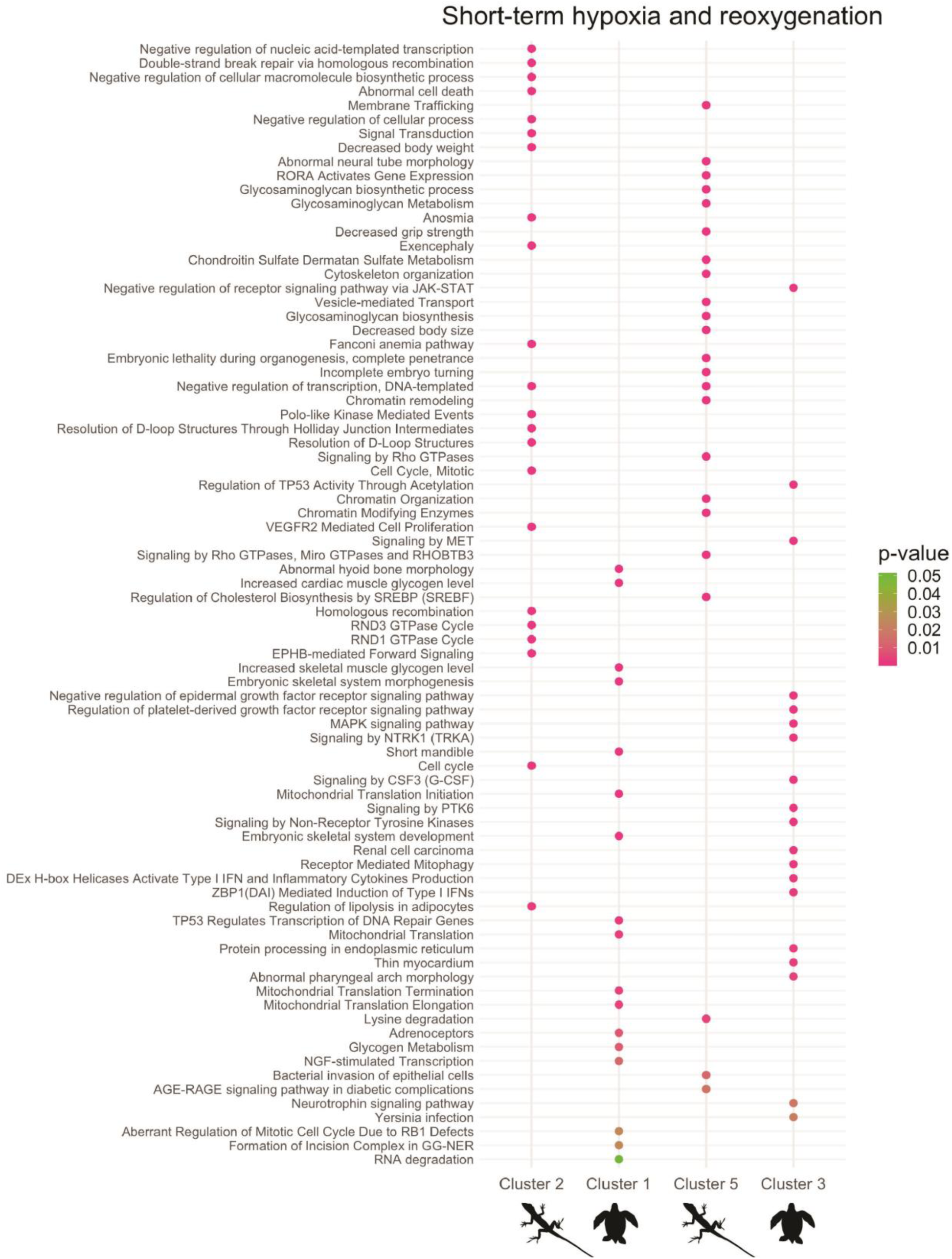
Complete list of enriched pathways from clusters with similar gene expression trajectories during short-term hypoxia and reoxygenation in lizard and sea turtle cells. Cluster 2 (lizard) and Cluster 1 (sea turtle), as well as Cluster 5 (lizard) and Cluster 3 (sea turtle), have matching trajectories.

## Notes

### Competing Interest Statement

The authors have declared no competing interest.

